# Strong genotype by environment interactions in the rice Global MAGIC population across seedling stage drought

**DOI:** 10.1101/2022.02.07.479433

**Authors:** Brook T. Moyers, Amelia Henry, Chitra Raghavan, Hein Zaw, Annarita Marrano, Hei Leung, John McKay

## Abstract

Crop adaptation is required to sustainably increase the rate of yield gains to meet projected future needs under the challenging conditions of climate change and competition for resources. Future adaptation will likely need to harness both highly polygenic traits and genotype-by-environment interactions (GxE), the study of which is aided by complex recombinant populations. We used the diverse *Oryza sativa* Global Multiparent Advanced Generation InterCross (MAGIC) population to study the genetic architecture and contributions of seedling emergence, establishment, and development to yield components under dry-direct seeding and seedling stage drought across three growing seasons. Dry-direct seeding is an establishment practice that has the potential to reduce methane emissions, water use, and labor demands for rice farmers, but increases the possibility of early-season drought conditions. We found substantial evidence for large roles of environmental variation and GxE in controlling trait variation. Maintenance of shoot growth during seedling stage drought was positively correlated with crown root number and both directly and indirectly influenced agronomic traits. Other than the major green revolution locus *sd1*, most allelic effects were conditionally neutral (affecting trait values in some environments but not others) and most alleles had their strongest effect in an environmental extreme. This discovery is both a challenge and a potential opportunity: with variable genetic architecture, selection in any one environment may not result in stable trait gains across environments. However, conditionally neutral GxE is a potential route to sustainable yield stability through allele pyramiding.

## Introduction

The green revolution of the 1960s jump-started an era of deliberate, intensive breeding for high input crop management, leading to global yields increasing linearly in the decades following. Unfortunately, this steady rate of yield increase is insufficient to meet projected future needs. Additionally, high inputs of water and fertilizer create greenhouse gas emissions and runoff that are deleterious to local and global ecosystems. In short, current crop production methods are unsustainable, and contribute to and are impacted by climate change (Tilman *et al*. 2011; Ray *et al*. 2013). The goal for researchers, breeders, and farmers is therefore to find new ways of increasing yields quickly, but with lower inputs. This need is especially pressing in rice (*Oryza sativa)*, which provides 20% of human calories and is the primary calorie source for most of the world’s poorest people (FAOSTAT 2011). Global rice production is a major source of methane emissions, contributing to accelerating climate change (Neue 1993; Wassmann *et al*. 2000). Rice mega-varieties developed since the green revolution are adapted to high inputs of water and fertilizer, and tend to perform poorly under less optimal growth conditions (Dobermann *et al*. 2002; Ismail and Atlin 2019). Sustainable yield growth in rice therefore requires both alternative production methods and new, robust varieties that yield well in more challenging conditions.

Transplanted, irrigated lowland production is the highest yielding rice cropping system (e.g. Devkota *et al*. 2020)), but is highly resource intensive and the major source of methane emissions in rice production (Wassmann *et al*. 2000). It is also not feasible in many regions. In contrast, establishment of rice by dry-direct seeding, a common practice in upland rice and most non-rice systems, holds promise for increased sustainability. Dry-direct seeding allows earlier and faster crop establishment with reduced labor, water, and greenhouse gas emissions (Pandey *et al*. 2002; Kumar and Ladha 2011). Transition to dry-direct seeding could potentially benefit rice farmers in many regions, including Eastern India, Central China, and the Mekong region (Fukai and Ouk 2013; Wang *et al*. 2017; Balwinder-Singh *et al*. 2019). To create new varieties optimized for dry-direct seeding, breeders must target germination traits and rapid early establishment to address challenges with weed competition and development of a good crop stand (Pandey *et al*. 2002; Kumar and Ladha 2011). With the absence of soil puddling, water losses due to percolation are greater in soils planted by direct seeding, making them prone to drought stress especially at the seedling stage (Haefele *et al*. 2016). Development of new varieties with both adaptation to dry-direct seeding and improved growth under early drought stress is key to widespread implementation of dry-direct seeding with sustainable yields.

Alleles with large, consistent effects, such as the green revolution allele *semidwarf-1* (*sd1*), are ideal for breeding in many ways: relatively simple to introgress, easy to monitor during marker-assisted selection, and complex effects across environments are not a concern. Such alleles are unfortunately rare, and breeders are left with the challenge of achieving yield and other trait stability in the face of genotype by environment (GxE) and epistatic interactions (Becker and Leon 1988). Alleles that can enable trait stability even with GxE are those with conditionally neutral effects, where an allele is beneficial in some environments but has minimal effect in others. Alleles with antagonistic effects, on the other hand, are beneficial in some environments and deleterious in others, resulting in environmental trade-offs. These concepts are central to understanding the genetics of adaptation in wild species (Anderson *et al*. 2011), and have great potential for application in crop adaptation.

Distinguishing between classes of allelic effect requires that recombinant populations be grown across multiple environments. One such diverse highly recombinant population design is MAGIC, or **M**ulti-parent **A**dvanced **G**eneration **I**nter**C**ross, which elaborates on traditional bi-parental crossing designs by increasing the number of recombining parent lines. MAGIC population designs advance lines through single seed descent until the lines are nearly completely homozygous, enabling experimental replication within environments and study of GxE. Quantitative genetic analyses in MAGIC populations combine the higher statistical power of recombinant populations with the higher resolution of association panels (Meng *et al*. 2016b). In addition, MAGIC lines represent novel allelic combinations that can expose novel epistatic interactions and novel trait values (Leung *et al*. 2015). In rice, researchers have used MAGIC populations within and across the *indica* and *japonica* subspecies to uncover the genetic bases of abiotic and biotic stress tolerances, grain quality, and a number of agronomic and yield component traits under transplanted lowland conditions (Meng *et al*. 2016b, 2016a; Raghavan *et al*. 2017; Bossa-Castro *et al*. 2018; Ogawa *et al*. 2018b, 2018a; Descalsota *et al*. 2018; Zaw *et al*. 2019). To date, few studies have used complex or diverse populations for genetic discovery and breeding in dry-direct seeded rice and all of these have focused on grain traits under well-irrigated conditions (Sandhu *et al*. 2019b; Subedi *et al*. 2019).

Given that drought often occurs with dry-direct seeding crop establishment, understanding how rice plants respond to this combination of conditions is important. To meet this aim, we grew the rice Global MAGIC population (Bandillo *et al*. 2013) under seedling stage drought stress across three growing seasons. Here we dissect the genetic and environmental contributors to seedling establishment, seedling stage drought response, and traits measured at reproductive stage and maturity. We evaluate support for three non-mutually exclusive hypotheses about the adaptive value of specific traits for dry-direct seeding: (1) early emergence contributes to increased yield under dry-direct seeding, (2) increased mesocotyl length creates a better distributed root system that improves plant growth, and (3) maintenance of seedling growth during drought is mediated by below-ground architecture and contributes to agronomic traits. We examine how genetic architectures vary across environments and explore how conditionally neutral allelic effects could be used in future rice breeding efforts.

## Materials and Methods

### Rice Global MAGIC population

The rice Global MAGIC population includes 1200 recombinant lines advanced from intercrosses between two eight-parent MAGIC populations meant to represent each of the two major Asian rice (*Oryza sativa*) subspecies *indica* and *japonica*, with one parent from the circum-Basmati or aromatic rice subpopulation (Bandillo *et al*. 2013; Wang *et al*. 2018; **Table S1**). The 16 parent lines represent a diverse set of globally popular varieties with desirable agronomic traits, with the dual goals of enabling fine-scale genomic mapping of phenotypes and generating breeding-ready germplasm with recombinant or transgressive trait values. Of the MAGIC populations developed at the International Rice Research Institute, the Global MAGIC is the only one with sixteen parents drawn from three *O. sativa* sub-populations so is likely the most genetically and phenotypically diverse.

### Phenotyping

We evaluated 568 Global MAGIC lines in four dry-direct seeded experiments at the International Rice Research Institute (IRRI; 14° 10’ N, 121° 15’ E). We use the term ‘environment’ to refer to each of these four experiments. We planted in three of these environments from 2014 to 2015 in the dry season (DS) and wet season (WS) under varying seedling stage drought conditions (2014DS: S5 generation, 2015DS: S6, and 2015WS: S7) and one environment under flooded, well-watered (ww) conditions (2015DS-ww: S6 generation). The MAGIC lines grown in each season (2014DS, 2015DS, 2015WS) mostly overlapped but were not identical due to availability of seed and plot space, with 437 MAGIC lines in all environments, 78 in three environments, 11 in two environments, and 42 in only one environment (**Table S2**). We planted in augmented block designs with the 16 parents, IRRI standard checks, and in 2015DS/2015DS-ww 10 promising MAGIC lines replicated 3–6 times and the remaining MAGIC lines unreplicated.

We hand-sowed in two of the environments (2014DS and 2015WS) by placing 3–4 seeds at depths of 2–3 cm every 5 cm in plots of two 2 m rows with 25 cm row spacing under a rolling rainout shelter. We mechanically sowed in the other two environments (2015DS and 2015DS-ww), placing one seed at a depth of 1–2 cm every 5.4 cm in plots of six 1.6 m rows with 20 cm row spacing in an open field, and planted the ww treatment on a lower terrace than the drought stress treatment. We sowed into dry, rotovated soil, and established plant growth by sprinkler irrigation 2–3 times per week or by moving the rainout shelter to allow rainfall to reach the plot. In 2015DS and 2015DS-ww, variable establishment led to gaps in the plant stands of some plots, which we filled with a visually differentiable purple-leafed genotype to avoid variation in neighbor effects.

In the drought stress environments, we stopped irrigation at 12 days after sowing (DAS) in 2014DS and 2015WS, and at 34 DAS in 2015DS due to slower establishment. We monitored the level of drought stress using tensiometers installed at 30 cm depth (**Figure S1**). Following the seedling stage drought treatments, we resumed irrigation at 70 DAS in 2014DS, 45 DAS in 2015DS, and 27 DAS in 2015WS based on the progression of the seedling stage drought in each environment. We maintained the 2014DS and 2015WS environments well-watered following the seedling stage drought stress treatments, and intermittently resumed the stress treatment in 2015DS with additional irrigations at 77 and 90 DAS. The 2014DS and 2015WS environments were not affected by rainfall during the drought stress treatment due to cover by the rainout shelter, and 2015DS received in total 117 mm of rainfall following the initiation of the drought stress treatment. We flooded the 2015DS-ww environment at 34 DAS and maintained it flooded throughout the season.

We collected measurements on both seedling and reproductive stage traits. In 2014DS and 2015WS, we recorded time to first emergence as the DAS that the first coleoptiles emerged in each plot, and time to full emergence as the DAS that the entire length of the planted rows had emerged. We calculated establishment time as the number of days between first and full emergence. We did not collect data on these traits during the 2015DS due to variable establishment after mechanical sowing. Immediately before rewatering each drought stress environment, we sampled three plants per plot for seedling-stage shoot and crown root measurements (at 70 DAS in 2014DS, 45 DAS in 2015DS and 2015DS-ww, and 27 DAS in 2015WS). The mean soil water potential at a depth of 30 cm on sampling dates were −52 kPa in 2014DS, −23.4 kPa in 2015DS, and −12 kPa in 2015WS (**Figure S1**). We excavated plants manually using a trowel from a depth of about 10 cm. We detached and dried shoots to determine shoot dry weight and washed and stored root crowns in 25% ethanol until root crown measurements were conducted. We measured mesocotyl length and axial root numbers for each of three classes: crown roots, adventitious (mesocotyl-borne) roots, and seminal roots. Adventitious roots were rare and therefore this trait was not included in our analysis. We censused plots three times weekly and recorded the number of days to flowering as the DAS at which 50% of each plot flowered. At flowering we also measured and calculated mean plant height from three plants in each plot. At maturity we harvested grain yield and straw biomass from areas of 0.5 m^2^ in 2014DS and ~0.23 m^2^ (20 plants) in 2015DS and 2015DS-ww. We did not measure mean height, grain yield, or straw biomass in 2015WS due to poor plant growth following seedling stage.

### Phenotypic analyses

To adjust non-replicated plots for block-level environmental variance, we estimated least squares means for each trait of each genotype in each environment using the *lme4* R package version 1.1-21 (Core Team and Others 2013; Bates *et al*. 2015) in the linear model y = μ + G + B + ε, where y is the plot value, μ is the overall mean, G is the random genotypic effect, B the block effect, and ε are the residuals. We scaled grain yield and straw biomass to grams per square meter. We calculated harvest index as the ratio of grain mass to total above-ground plant biomass (straw plus grain mass). We calculated Drought Response Index (DRI) for the 2015 dry season drought environment using the time to flowering and grain yield data from 2015DS and 2015DS-ww following Bidinger *et al*. (1987). We calculated yield stability indices using the AMMI function of the R package *agricolae* version 1.3-1 (De Mendiburu and Simon 2015), and tested for differences using permutation testing with 1000 random subsamples of the same length as our observed set. We identified transgressive segregants in the MAGIC lines as lines with trait values outside of the range of the first quantile minus 1.5 * IQR to the third quantile plus 1.5 * IQR of the distribution of parent line trait values for that environment, and tested for increased variance in the MAGIC lines relative to the parent lines using one-tailed Ansari-Bradley tests (which do not assume normally-distributed data or equal sample sizes).

To examine patterns of phenotypic trait variance, we conducted a principal components analysis for all traits of each Global MAGIC line in the four environments using the *FactoMineR* R package version 1.42 (Lê *et al*. 2008). As this analysis does not handle missing data well, we imputed missing individual values using the imputePCA() function from the *missMDA* R package version 1.14, with the number of principal components (PC) used in imputation estimated by the estim_ncpPCA() function (Josse *et al*. 2016). We examined all PCs in the final PCA with eigenvalues greater than 1. We tested for phenotypic differences between treatments (drought stress and well-watered) and growing seasons (dry and wet) using ANOVA of linear models with the formula PC_X ~ treatment + season. We calculated genetic correlations among traits as the correlation between genotypic values for each pair of traits, with genotypic values estimated as the mean trait value for each genotype across all four environments.

To explore our three hypotheses of adaptation to dry-direct seeding we modeled direct and indirect effects of earlier developmental traits on later traits using structural equation modeling (SEM) as implemented in the R package lavaan (version 0.6–8, Rosseel 2012). For each environment, our initial model consisted of each developmentally earlier trait as a direct predictor of each later trait, with traits from similar developmental periods co-varying without implied directionality of effect. Specifically, first emergence, full emergence, and mesocotyl length co-vary and predict seedling biomass, crown root number, and seminal root number, which co-vary, and all together predict days to flower and height, which co-vary and all together predict straw biomass, which all together predict grain yield. The models did not include composite traits calculated from traits already in the model, namely establishment time, total axial roots, and harvest index, as these would introduce issues with collinearity. Where within-environment trait variances were three or more orders of magnitude apart, we scaled trait values to avoid traits with large variances having disproportionate effects on model estimation. We fit these models to the raw per plot trait data with a robust maximum likelihood estimation (estimator = “MLR” in lavaan::sem) that can handle missing and non-normally distributed data, and used case-wise maximum likelihood estimation (missing = “ML” in lavaan::sem). Our initial models were saturated (i.e. no remaining degrees of freedom to examine model fit), so we dropped non-exogenous regression predictors that were not significant (p < 0.05) in any environment from our final models: crown roots never affected days to flower, seminal roots never affected height or straw biomass, and seedling biomass never affected grain yield. We also fit models for the 2014DS data without emergence traits (2014DS-noem, to directly compare against both 2015DS and 2015DS-ww), and without agronomic traits (2014DS-early, to compare against 2015WS). All final models showed good fit to the data as estimated by non-significant χ2 tests, robust Root Mean Square Error of Approximation (RMSEA) < 0.02 and robust Comparative Fit Index (CFI) > 0.99 (**Table S3**). In describing these models we primarily report standardized parameters where both latent and observed variables have a variance of one (similar to standardized regression coefficients).

### Genotyping

1184 recombinant inbred lines of the S6 generation of the Global MAGIC population were genotyped from DNA extracted from leaf tissue using Genotyping-by-Sequencing (GBS, with the enzyme *ApeKI*) across 14 lanes of 96-sample multiplexed sequencing (100 bp single-end Illumina, Phred+33 fastq format) as reported in Zaw *et al*. (2019). This sequencing includes all RILs that were grown as part of our field experiments. Each Global MAGIC S6 line was represented by a single individual extracted as a DNA template, while the 16 parent lines were represented by six biological replicates each. We used our own pipeline to identify genetic variants from these data aligned to two genomes: the gold standard Nipponbare (a japonica variety, assembly IRGSP-1.0/MSU7; Kawahara *et al*. 2013) and the highest quality indica genome, 93-11 (assembly ASM465v1; Zhao *et al*. 2004).

We used the TASSEL 5.0 GBS v2 pipeline (Glaubitz *et al*. 2014) to identify single nucleotide polymorphisms (SNPs). We first created a database of ‘tags’ using the GBSSeqToTagDBPlugin (minimum kmer count 100, kmer length 75, minimum kmer length 20, minimum quality score 20, maximum kmers in database 100M). The resulting database contained 917,254 tags in 1200 taxa (collapsing biological replicates for parents into a single taxon; separately we also called the biological replicates independently for filtering purposes). We used the TagExportToFastqPlugin to generate a fastq-formatted file containing all tag sequences for alignment. We duplicated this reference-free database for use against each reference genome, and all further bioinformatic steps were performed against each reference. The two genomes are largely collinear (**Figure S2**), and where we present analysis of only one alignment it is from the Nipponbare reference, with the highest-quality annotation and other resources.

We used Bowtie2-2.2.6 to align tags to each reference (with --sensitive preset mode; **Table S4;** Langmead *et al*. 2019). Approximately 9% of tags did not align to each reference genome, and 79% of those that did not align to one reference also did not align to the other reference (**Table S4**). We used the SAMToGBSdbPlugin to store alignment positions for each tag in the database. We used the DiscoverySNPCallerPlugin to identify variants (SNPs and indels) from aligned tags using the database (including more than 2 alleles/site, minimum locus coverage 0.1, minimum minor allele frequency 0.01). We then used the SNPQualityProfilerPlugin to score all discovered variants for coverage, depth, and genotypic statistics. We exported genotype data for all samples in VCF format using the ProductionSNPCallerPluginV2.

We filtered the resulting variants using statistics from a genotype dataset of the six biological replicates of the 16 parent lines and custom bash shell scripts. We filtered for variants in this dataset with > 50% coverage, no heterozygous calls, bi-allelic sites (including single indels), and consistent genotype calls for all replicates of a line. We then filtered for sites at which at least 15 of the 16 parent lines had genotype calls (when replicates were collapsed into a single taxon). Because heterozygotes were extremely rare in the MAGIC lines (0.009% in the Nipponbare-aligned dataset) and therefore could have disproportionate effects on our genome-wide associations, we masked heterozygote genotypes for all MAGIC lines. The phenotypic data was collected from the S5 and S7 generations as well as the genotyped S6 generation, so masking heterozygotes also helped to reduce the likelihood of unaccounted for genotype differences across generations. We excluded the eight MAGIC lines with > 90% missing data (all other MAGIC lines mean missing genotypes ~0.2 for both genomes, **Table S4**).

### Genome-wide Association

We looked for statistical associations between traits and genetic markers using a general linear model (GLM) approach as implemented in TASSEL 5 (Glaubitz *et al*. 2014). GLMs take as input genotype and phenotype matrices, but do not take kinship or relatedness among taxa into account. The Global MAGIC population should, by design, exhibit little to no population structure (Bandillo *et al*. 2013). We confirmed this by examining a kinship matrix among all 1176 post-filtering genotyped Global MAGIC S6 lines, estimated using the Centered Identity-by-State method (Endelman and Jannink 2012). Population structure is essentially flat (**Figure S3**; mean kinship = −0.001 ± 0.106 SD), which allowed us to examine genome-wide associations using simple and powerful GLMs. We ran GLMs (with structure y = μ + xβ + ε, where y is the phenotype, x is the additive genotype, μ is the mean phenotype of one genotype, β is the effect of each copy of the alternate allele, and ε are the residuals) for all traits within each environment and corrected for multiple testing using *p*-values from 1,000 permutations. We used an arbitrary threshold of permuted *p* < 0.1 to filter putative associations for further analysis, for a filtered set with a maximum unpermuted *p* of 0.00004. This relatively relaxed threshold allowed us to explore two traits (seminal roots and seedling biomass) and one environment (2015WS) with only marginal genetic associations, but did not obscure associations with higher confidence. We defined association peaks as sets of significantly associated markers within an LD window of 2 Mb. We explored putative candidate genes and links to other QTL using community databases: the Rice Annotation Project (Sakai *et al*. 2013), Q-TARO (Yonemaru *et al*. 2010), and Gramene (Ware *et al*. 2002).

We estimated narrow sense heritability (*h*^2^) for each trait in each environment as the ratio of additive genetic variation to the sum of additive genetic variation and residual variation, as calculated in TASSEL 5 using mixed linear models of genotype, phenotype, and the kinship matrix described above (which provides a good estimate of additive genetic variance; (Endelman and Jannink 2012). We also estimated *h*^2^ for each trait in each environment using variance components extracted from ANOVA of the replicated parent and check lines in the augmented designs (Conner and Hartl 2004).

### Data Availability

The raw sequence data are available at https://doi.org/10.7910/DVN/CANBL7. Our phenotypic and genotypic datasets, along with the scripts used for statistical analysis and to generate tables and figures, are available on the IRRI dataverse: [link to be updated upon acceptance].

## Results

### The rice Global MAGIC population harbors extensive phenotypic variation

In all four environments, Global MAGIC lines showed transgressive values for almost all traits (**Figure 1; Figure S4**). The traits with the strongest evidence for increased variance in the MAGIC lines include crown root number, total axial root number, and seedling biomass in both dry season drought environments, seminal roots and mesocotyl length in 2014DS, grain yield in 2015DS, straw biomass in 2015DS-ww, and days to full emergence in 2015WS (one-tailed Ansari-Bradley tests *p* < 0.05; **Table S5**). Substantially more MAGIC lines had transgressive values for crown root number and seedling biomass in the dry season drought environments (2014DS and 2015DS) relative to the well-watered environment (2015DS-ww), while the inverse is true for straw mass and grain yield (**Table S5**).

**Figure 1.**
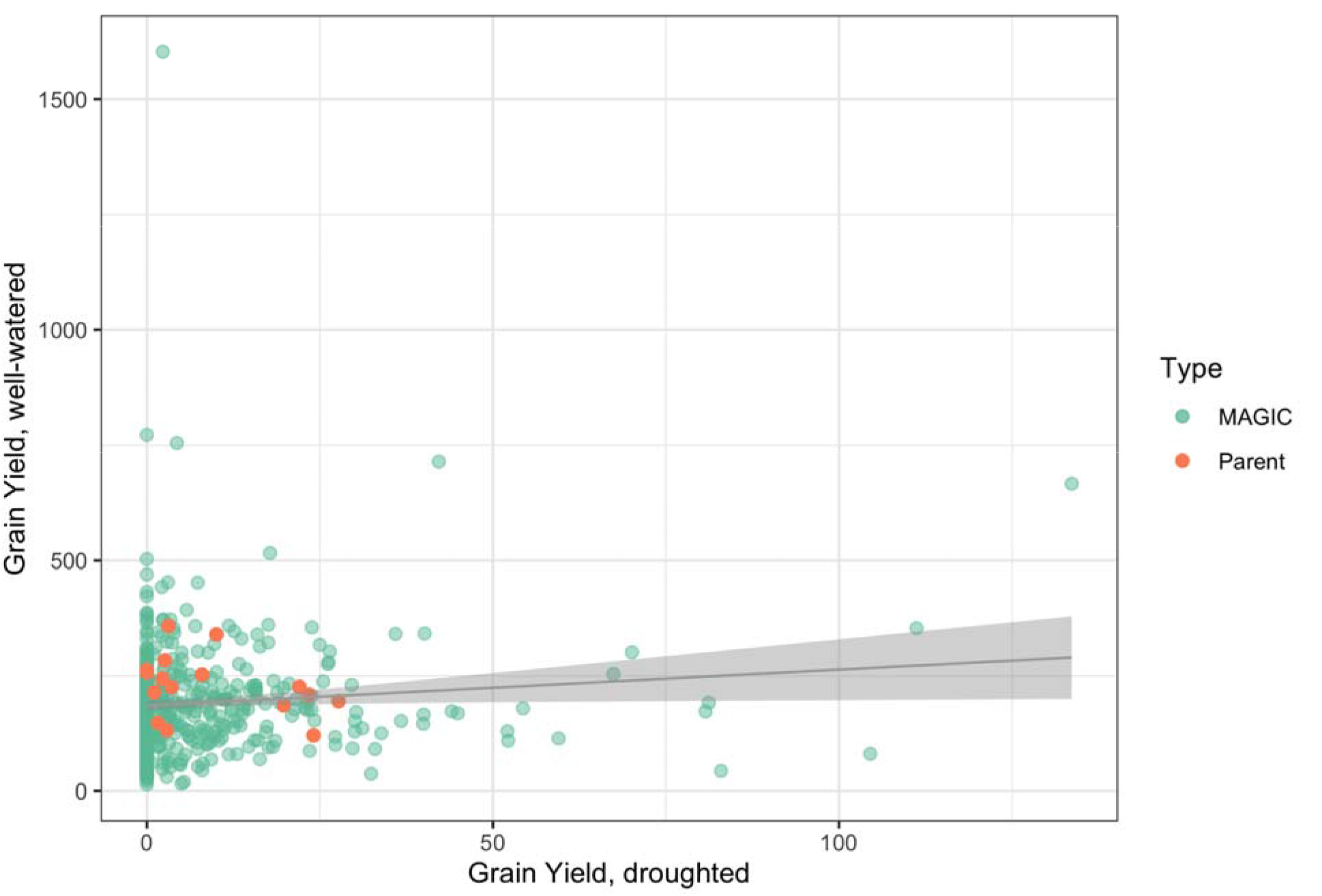
Grain yield (g m^-2^) for rice Global MAGIC S6 lines versus 16 parent lines under seedling-stage drought stress (x-axis) and under well-watered conditions (y-axis) during the 2015 dry season. Values are least-squared means for genotype, and the line depicts the linear regression ± standard error.

Trait heritabilities *(h^2^*) varied by trait, environment, and method of estimation (**Table 1**). We estimated heritability from the repeated lines in our augmented design using ANOVA variance components. The repeated lines are largely the parent lines, and these estimates of heritability were relatively high (mean 0.25, max. 0.9). In contrast, heritabilities estimated within the unreplicated MAGIC lines using relatedness were smaller (mean 0.10, max. 0.35) and uncorrelated with the repeated lines heritabilities (Pearson’s product moment correlation, *t* = - 0.092, df = 39, *p* = 0.927). This observation suggests that additive genetic variance played a smaller role in shaping trait variance in our MAGIC lines than would be predicted by the parent lines, with a likely larger role of environmental variance, epistatic variance, or genotype by environment interactions (GxE).

**Table 1.** Sample sizes and heritabilities for each trait in each environment, estimated two ways: from the MAGIC lines using kinship and the repeated parent lines using ANOVA. The ANOVA also tests whether repeated lines genotype is a predictor of trait values in that environment, and only heritability for traits with *p* < 0.05 are shown. GWA hits are trait-associated genetic variants with permuted *p* < 0.1, with GWA peaks defined as in **Table S9**.

| Trait | Environment | N | F | df | p-value | $h^2$, repeated lines ANOVA | $h^2$, MAGIC kinship | GWA hits | GWA peaks |
| --- | --- | --- | --- | --- | --- | --- | --- | --- | --- |
| First Emergence | 2014DS | 538 | 1.1859 | 20 | 2.90E-01 | — | 0.28 | 0 | 0 |
| First Emergence | 2015WS | 457 | 1.1275 | 20 | 3.78E-01 | — | 0.03 | 0 | 0 |
| Full Emergence | 2014DS | 531 | 2.0038 | 20 | 1.65E-02 | 0.18 | 0.11 | 0 | 0 |
| Full Emergence | 2015WS | 441 | 0.4266 | 19 | 9.71E-01 | — | 0.08 | 0 | 0 |
| Establishment Time | 2014DS | 531 | 0.8408 | 15 | 6.30E-01 | — | 0.13 | 0 | 0 |
| Establishment Time | 2015WS | 440 | 0.9258 | 17 | 5.55E-01 | — | 0.19 | 0 | 0 |
| Crown Roots | 2014DS | 521 | 1.7577 | 20 | 4.26E-02 | 0.14 | 0.14 | 7 | 3 |
| Crown Roots | 2015DS | 475 | 2.3861 | 22 | 5.06E-03 | 0.30 | 0.08 | 0 | 0 |
| Crown Roots | 2015DS-ww | 412 | 1.2305 | 23 | 2.79E-01 | — | 0.09 | 1 | 1 |
| Crown Roots | 2015WS | 315 | 1.3355 | 17 | 2.95E-01 | — | 0.00 | 0 | 0 |
| Seminal Roots | 2014DS | 521 | 1.3452 | 20 | 1.79E-01 | — | 0.00 | 0 | 0 |
| Seminal Roots | 2015DS | 475 | 1.0769 | 22 | 3.99E-01 | — | 0.00 | 0 | 0 |
| Seminal Roots | 2015DS-ww | 412 | 0.7891 | 23 | 7.23E-01 | — | 0.13 | 0 | 0 |
| Seminal Roots | 2015WS | 314 | 2.8676 | 17 | 2.62E-02 | 0.51 | 0.00 | 2 | 1 |
| Total Axial Roots | 2014DS | 521 | 2.0466 | 20 | 1.41E-02 | 0.19 | 0.15 | 9 | 3 |
| Total Axial Roots | 2015DS | 475 | 2.3101 | 22 | 6.66E-03 | 0.28 | 0.08 | 0 | 0 |
| Total Axial Roots | 2015DS-ww | 412 | 1.2184 | 23 | 2.88E-01 | — | 0.09 | 1 | 1 |
| Total Axial Roots | 2015WS | 315 | 3.2682 | 17 | 1.51E-02 | 0.56 | 0.01 | 0 | 0 |
| Mesocotyl Length | 2014DS | 521 | 1.3187 | 20 | 1.95E-01 | — | 0.00 | 18 | 2 |
| Mesocotyl Length | 2015DS | 475 | 0.7832 | 22 | 7.31E-01 | — | 0.06 | 0 | 0 |
| Mesocotyl Length | 2015DS-ww | 412 | 1.0986 | 23 | 3.89E-01 | — | 0.35 | 32 | 5 |
| Mesocotyl Length | 2015WS | 315 | 2.5722 | 17 | 4.02E-02 | 0.47 | 0.06 | 1 | 1 |
| Seedling Biomass | 2014DS | 534 | 0.7589 | 20 | 7.52E-01 | — | 0.13 | 2 | 1 |
| Seedling Biomass | 2015DS | 475 | 1.0692 | 22 | 4.07E-01 | — | 0.08 | 0 | 0 |
| Seedling Biomass | 2015DS-ww | 412 | 0.7327 | 23 | 7.83E-01 | — | 0.07 | 0 | 0 |
| Seedling Biomass | 2015WS | 315 | 2.5814 | 17 | 3.97E-02 | 0.47 | 0.05 | 0 | 0 |
| Days to Flower | 2014DS | 507 | 39.4388 | 19 | 6.28E-30 | 0.90 | 0.07 | 0 | 0 |
| Days to Flower | 2015DS | 415 | 4.2571 | 22 | 2.16E-05 | 0.53 | 0.00 | 0 | 0 |
| Days to Flower | 2015DS-ww | 509 | 3.6563 | 22 | 6.79E-05 | 0.45 | 0.01 | 0 | 0 |
| Days to Flower | 2015WS | 236 | 1.5200 | 16 | 3.16E-01 | — | 0.11 | 0 | 0 |
| Height | 2014DS | 528 | 9.1698 | 20 | 9.71E-13 | 0.65 | 0.23 | 40 | 9 |
| Height | 2015DS | 522 | 4.1517 | 23 | 5.84E-06 | 0.48 | 0.10 | 15 | 1 |
| Height | 2015DS-ww | 517 | 2.0331 | 23 | 1.53E-02 | 0.23 | 0.29 | 29 | 1 |
| Straw Biomass | 2014DS | 526 | 4.7717 | 20 | 4.38E-07 | 0.46 | 0.11 | 8 | 1 |
| Straw Biomass | 2015DS | 517 | 0.9668 | 23 | 5.18E-01 | — | 0.01 | 0 | 0 |
| Straw Biomass | 2015DS-ww | 497 | 1.7620 | 22 | 5.02E-02 | 0.20 | 0.04 | 13 | 1 |
| Grain Yield | 2014DS | 531 | 15.0168 | 20 | 2.00E-18 | 0.76 | 0.08 | 2 | 1 |
| Grain Yield | 2015DS | 517 | 1.9855 | 23 | 1.87E-02 | 0.23 | 0.13 | 3 | 2 |
| Grain Yield | 2015DS-ww | 493 | 1.4790 | 22 | 1.27E-01 | — | 0.07 | 2 | 2 |
| Harvest Index | 2014DS | 523 | 17.2819 | 20 | 1.19E-19 | 0.79 | 0.19 | 2 | 1 |
| Harvest Index | 2015DS | 517 | 5.0073 | 23 | 3.86E-07 | 0.54 | 0.12 | 3 | 2 |
| Harvest Index | 2015DS-ww | 493 | 1.5120 | 22 | 1.14E-01 | — | 0.26 | 13 | 3 |
| Drought Response Index | 2015DS | 480 | 2.2968 | 20 | 9.25E-03 | 0.28 | 0.09 | 2 | 2 |

### Seedling stage drought has consistent phenotypic effects

We designed this study to capture early traits (emergence, establishment, root and shoot development following seedling stage drought) that we expected to consistently impact reproductive stage (agronomic) traits. We find a strong multi-trait response to drought: the first phenotypic principal component (PC1) explains 50.8% of individual trait variance and strongly differentiates the well-watered from drought-stressed treatments (*p <<* 0.0001) and between growing seasons (*p <<* 0.0001; full model *F*_2,2043_ = 2899,*p <<* 0.0001, adj. *R*^2^ = 0.739; **Figure 2**, **Table S6**). Emergence and establishment time, crown and total axial root number, seedling biomass, and most agronomic traits correlate strongly positively with PC1, while seminal root number and days to flowering correlate negatively (**Table S7**). The direction of phenotypic change was consistent across the drought environments despite differences in the timing and extent of drought conditions and in seasonality (**Figure S1**). The strongest phenotypic correlations were within each environment (triangles on the diagonal in **Figure 3**), especially among traits collected at the same developmental stage (*i.e*. within establishment traits, seedling traits, or reproductive traits).

**Figure 2.**
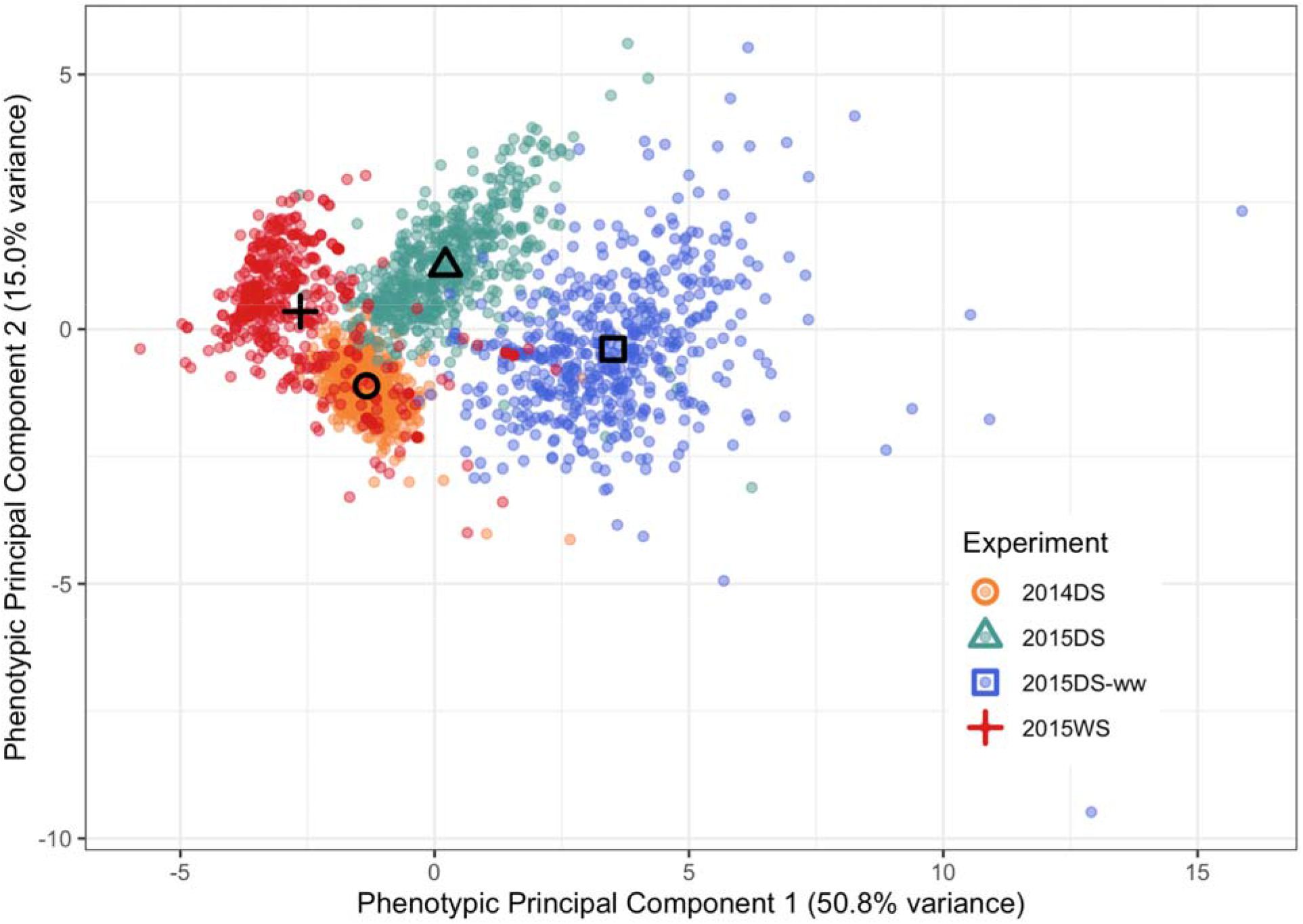
The first two phenotypic principal components of all traits in the four environments. Each point represents the phenotypic values for a Global MAGIC line in one environment, colored by environment. Environmental mean values are represented by shapes. Eigenvalues are in **Table S6** and trait correlations with each PC are in **Table S7**.

**Figure 3.**
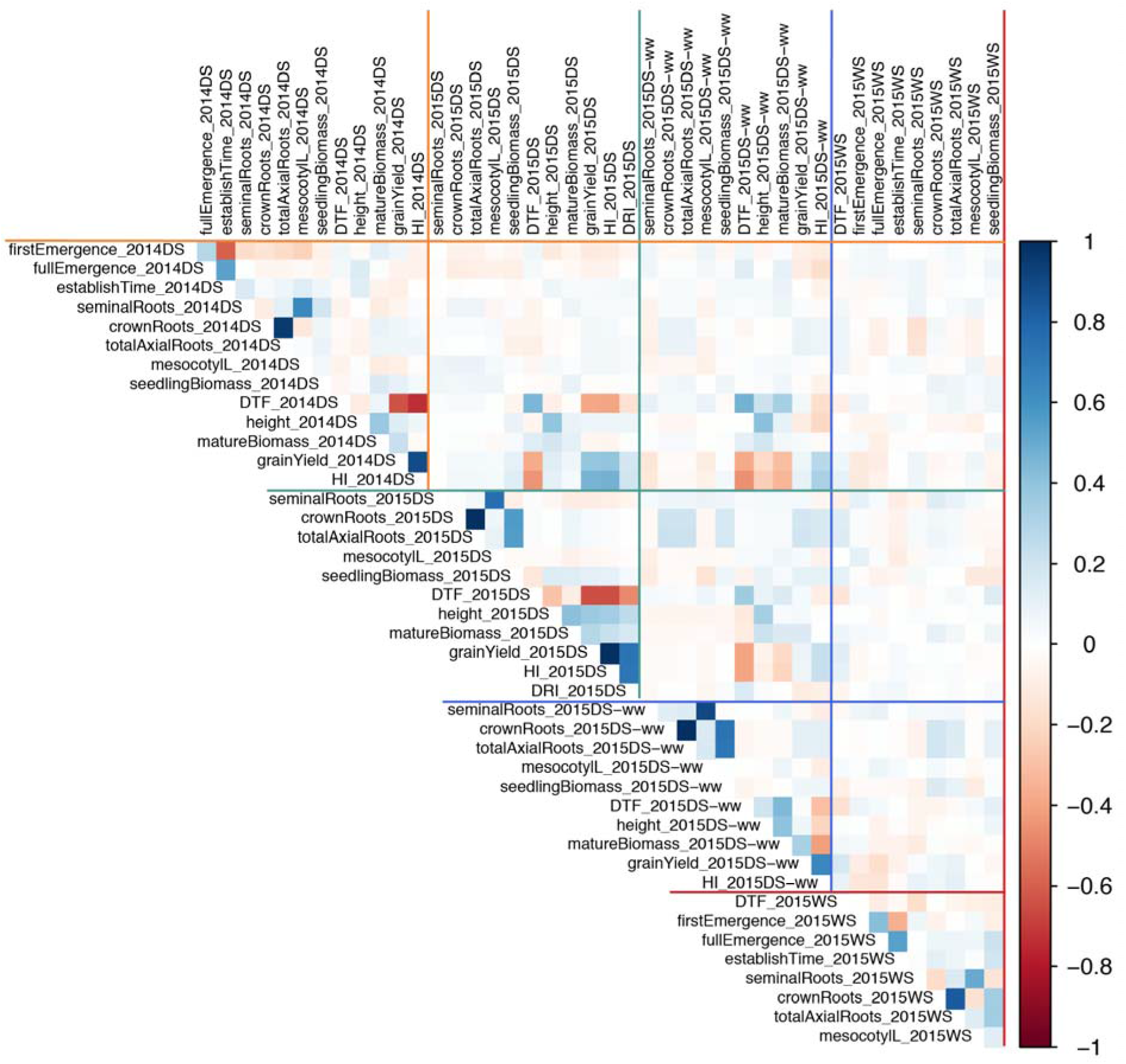
Pairwise phenotypic correlations (Spearman’s *rho*) for all traits measured in all experiments. Bluer squares indicate larger positive *rho* values while redder squares represent larger negative *rho* values, and traits are ordered by developmental stage within experiments separated by lines.

### Phenotypic patterns within and across environments

Grain yields in the 2015 dry season were positively correlated across the drought and well-watered environments, although the proportion of variance explained by yields from one environment to the other is low (*F*_1,499_ = 4.744, *p* = 0.02986, adj. *R*^2^ = 0.007433; **Figure 1**). Across environments (squares away from the diagonal in **Figure 3**), days to flower and height were each consistently positively correlated. Days to flower was also consistently negatively correlated with grain yield and harvest index across environments. We see few trait correlations between any of the dry season environments and the wet season environment, where we were not able to measure agronomic traits (rightmost column of square in **Figure 3**).

Season is a significant predictor of individual variance along the first four phenotypic principal components that cumulatively explain ~87% of individual variance (**Table S6**). Lines grown in 2015WS were quicker to establish than in 2014DS but seedlings were overall smaller than in either dry season drought environment. Although flowering times were generally positively correlated across environments (**Figure 3**), a group of 46 lines flowered very early in the wet season (< 50 days) and relatively late in the dry season experiments (mean DTF 101 in DS2014, 99 in DS2015, and 89 in DS2015-ww), indicating that day length sensitivity is segregating in this population (**Figure S5**). This group included two parents, IR-84196-32 (*i.e*. Sambha Mahsuri + *Sub1*) and WAB 56-125.

Some patterns in our structural equation models were consistent across environments. In both the 2014DS and 2015WS models, earlier first emergence strongly predicted earlier full emergence (**Figure 4; Table S8**). Crown root number and seedling biomass co-varied positively in every model, with standardized covariances from 0.14 in 2015WS to 0.74 in 2015DS-ww. The effect of mesocotyl length on seminal root number was the strongest positive predictor across all models except 2015WS, where it still had a substantial positive effect. Height was a consistent positive predictor of straw biomass in all DS environment models. In those same models, the effect of days to flower on grain yield was the strongest negative predictor, with standardized parameters of −0.63 in 2014DS, −0.37 in 2015 DS, and −0.20 in 2015DS-ww.

**Figure 4.**
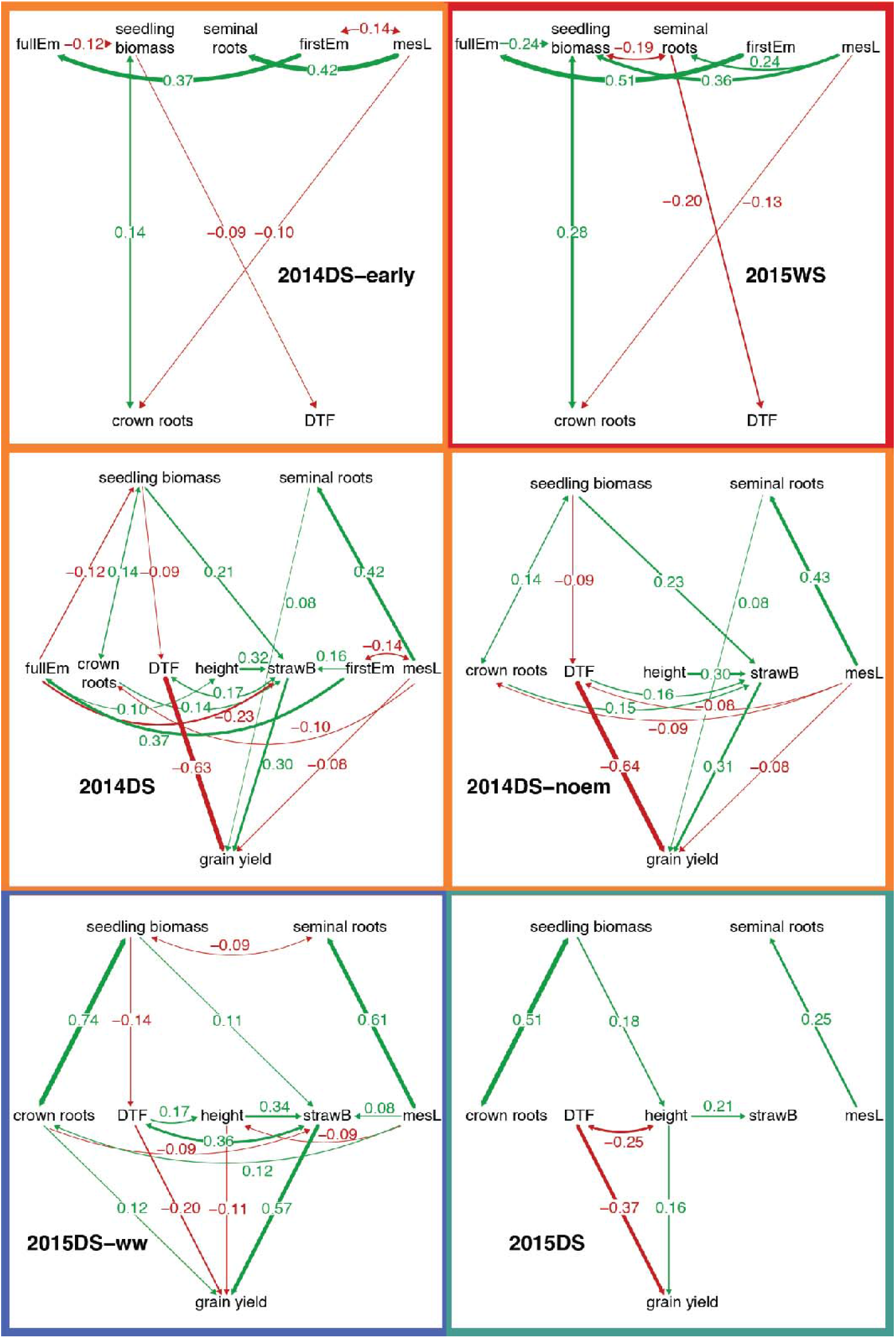
Structural equation models within each environment. We examined two additional models for 2014DS with only traits measured in 2015WS (2014DS-early) and 2015DS (2014DS-noem) for comparison. Traits are the model nodes (firstEm = first emergence, fullEm = full emergence, mesL = mesocotyl length, DTF = days to flower, and strawB = straw biomass, all others as written) connected by either two-directional arrows (covariances) or one-directional arrows (predictors). All trait relationships with *p* < 0.05 are shown. Arrows are colored by the direction of effect (red = negative, green = positive) and labeled with parameters standardized by setting all variable variances to one, which are also reflected in arrow width. See **Table S8** for additional model details.

Most other trait relationships varied across environments in our models. For example, seminal roots co-varied negatively with seedling biomass in 2015DS-ww and 2015WS but not 2014DS or 2015DS (**Figure 4; Table S8**). Seminal roots negatively predicted days to flower only in 2015WS and positively predicted grain yield only in 2014DS. Seedling biomass was a strong positive predictor of straw biomass and a negative predictor of days to flower only in 2014DS and 2015DS-ww. We observe the same pattern for straw biomass as a positive predictor of grain yield: seedling biomass indirectly increased grain yield through straw biomass in 2014DS and 2015DS-ww but had a different indirect effect through height in 2015DS. Height predicted grain yield positively in 2015DS, negatively in 2015DS-ww, and was not a significant predictor in 2014DS. Together, this variation in model structure provides additional support that environmental variance or GxE were important contributors to phenotypic variance.

### Genome-wide associations are environment-dependent

After filtering we have 7795 variable markers aligned against the Nipponbare genome and 7607 aligned against 93-11 (**Table S4**). The Nipponbare-aligned sequences from which these SNPs are derived overlap with 82% of the 93-11 aligned sequences, with most but not all sequences aligned colinearly (**Figure S2**). The SNPs are genome-wide but somewhat clustered, comparing the mean to median inter-SNP distances of 83 kb to 17 kb for Nipponbare and 86 kb to 19 kb for 93-11, respectively. Read coverage per genotyped site for each line is ~30X for parents and 5–6X for the 1176 MAGIC lines (**Table S4**). Allele frequencies in the subset of lines used for phenotyping are highly correlated with those in the full S6 population (**Figure S6**).

We observe strong genotype-phenotype associations (permuted *p* < 0.05) for all agronomic traits (except days to flower) and for crown root number, total axial root number, and mesocotyl length (**Figure 5, Table S9**). We also see a few marginal associations (0.05 < permuted *p* < 0.1) for seminal root number and seedling biomass. Most of the associations are observed in the three dry season environments, with only two weak associations in 2015WS, for mesocotyl length and seminal roots. We see consistent effects across environments at only one GWA peak: for height on chromosome 1 around 38–39 Mb (**Figure 6; Table S9**). All other associations are observed in only one or two environments, which could be due to greater environmental variance than genetic variance on that trait in the other environments. This potentially explains why we see few associations in 2015WS, but is unlikely to be the primary driver of trait variation in the dry season environments, where we see different genomic regions strongly associated with the same trait. A likely alternative is GxE, where allelic effects depend on environmental context.

**Figure 5.**
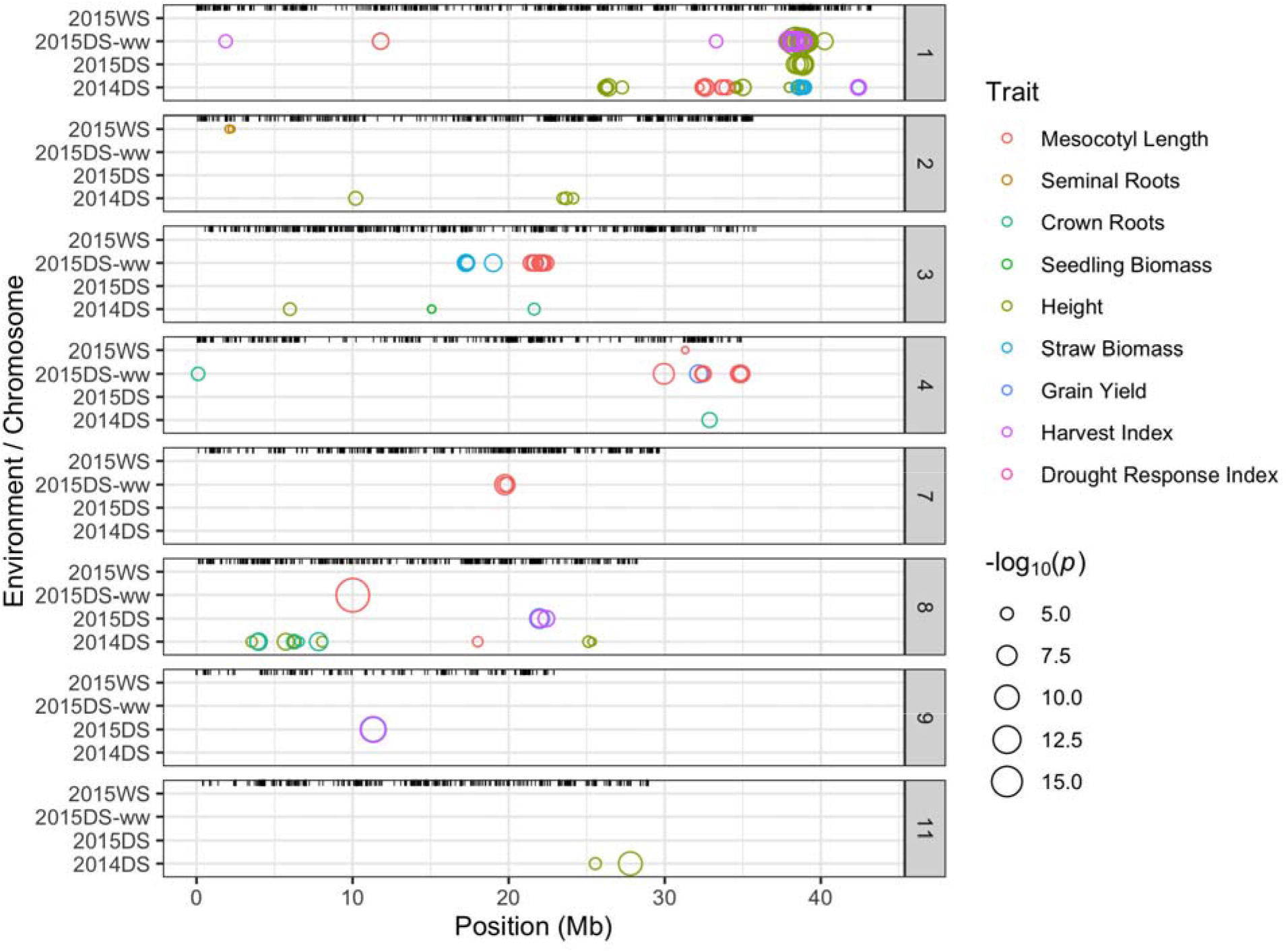
Most marker-trait associations are environment-specific, with the exception of sites linked to *SD1* (near Chr 1, Mb 38.4). Genome-wide associations with permuted *p* < 0.1 plotted as open circles colored by the associated trait in each environment (y-axis) along chromosomes (x-axis for position, chromosomes on vertical panels). Circle size is proportional to - log10(unpermuted *p*). Location of all markers is shown in a rug at the top of each panel. Only chromosomes and traits with genome-wide associations in at least one environment are shown. Total axial roots not shown: associations are identical to crown roots.

**Figure 6.**
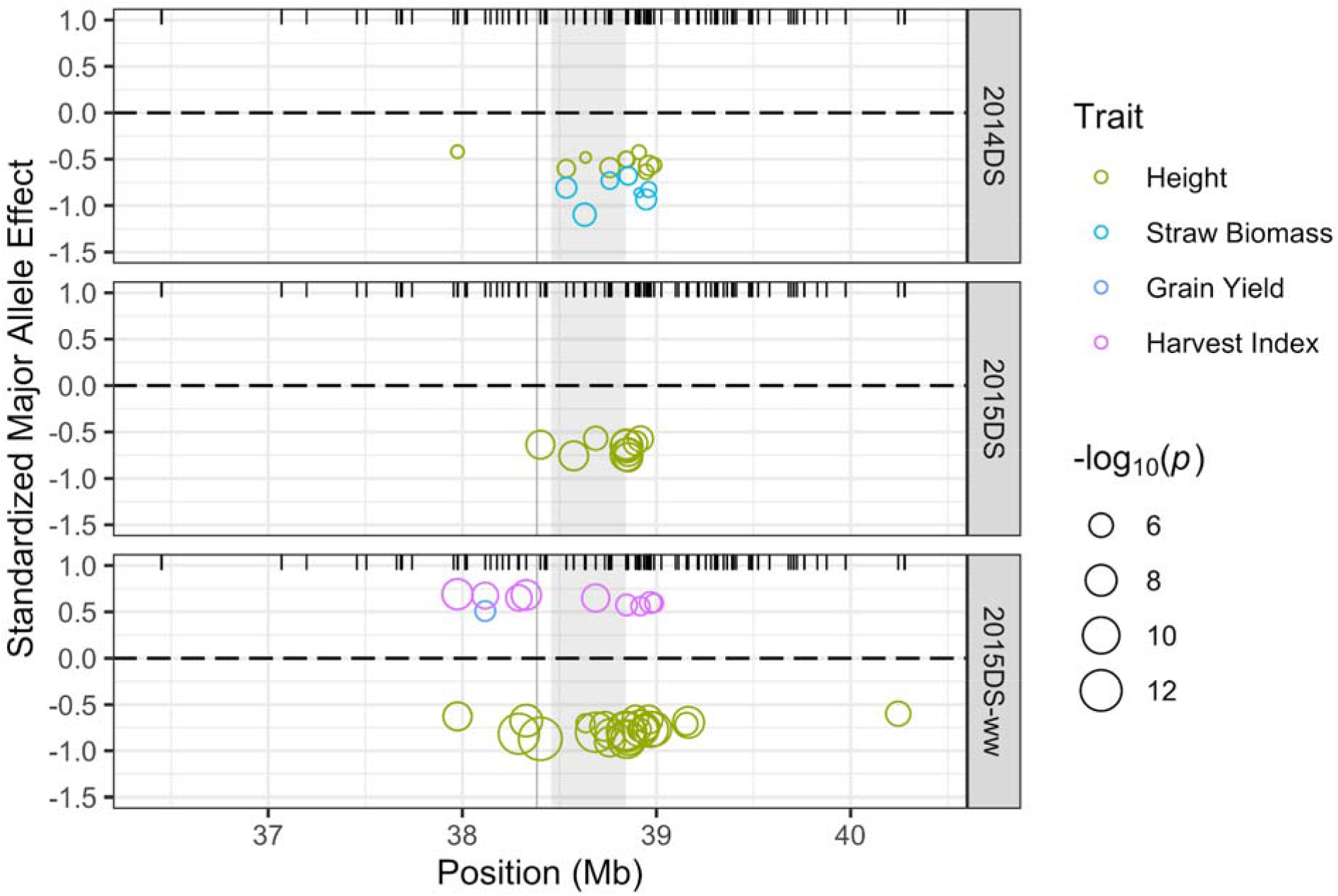
Major allele *sd1* constitutively reduces plant height but only increases grain yield and harvest index in the well-watered environment. Standardized effects of the major allele (y-axis) for genome-wide associations with permuted *p* > 0.1 plotted as open circles colored by the associated trait along rice chromosome 1 (x-axis) near *sd1* (vertical grey line) and *qDTY1.1* (grey shaded area). Circle size is proportional to −log10(unpermuted *p*). Location of all sites shown in a rug at the top of each panel.

We assessed GxE allelic effects using trait mean in each environment as our best proxy for environmental differences affecting that trait. We generally see larger absolute effects when mean trait values are higher (**Figure S7**), as is expected if variances are also higher. Comparing absolute against standardized allelic effects reveals that sometimes the relative impact of an allele is larger in the more stressful environments (e.g. compare grain yield effects in **Figure 7** to **Figure S7**). For all traits except seminal root number, the well-watered environment had the highest mean trait value and one of the drought stressed environments had the lowest mean trait value: 2015DS for mesocotyl length, height, grain yield, and harvest index, 2014DS for crown root number and straw biomass, and 2015WS for seedling biomass (**Figure 7**). This pattern is additional evidence that each drought stressed environment had unique impacts across development, possibly through differences in drought intensity and duration (**Figure S1**). For seminal root number the drought-stressed environments had higher mean values than 2015DS-ww, with 2015WS at the high extreme.

**Figure 7.**
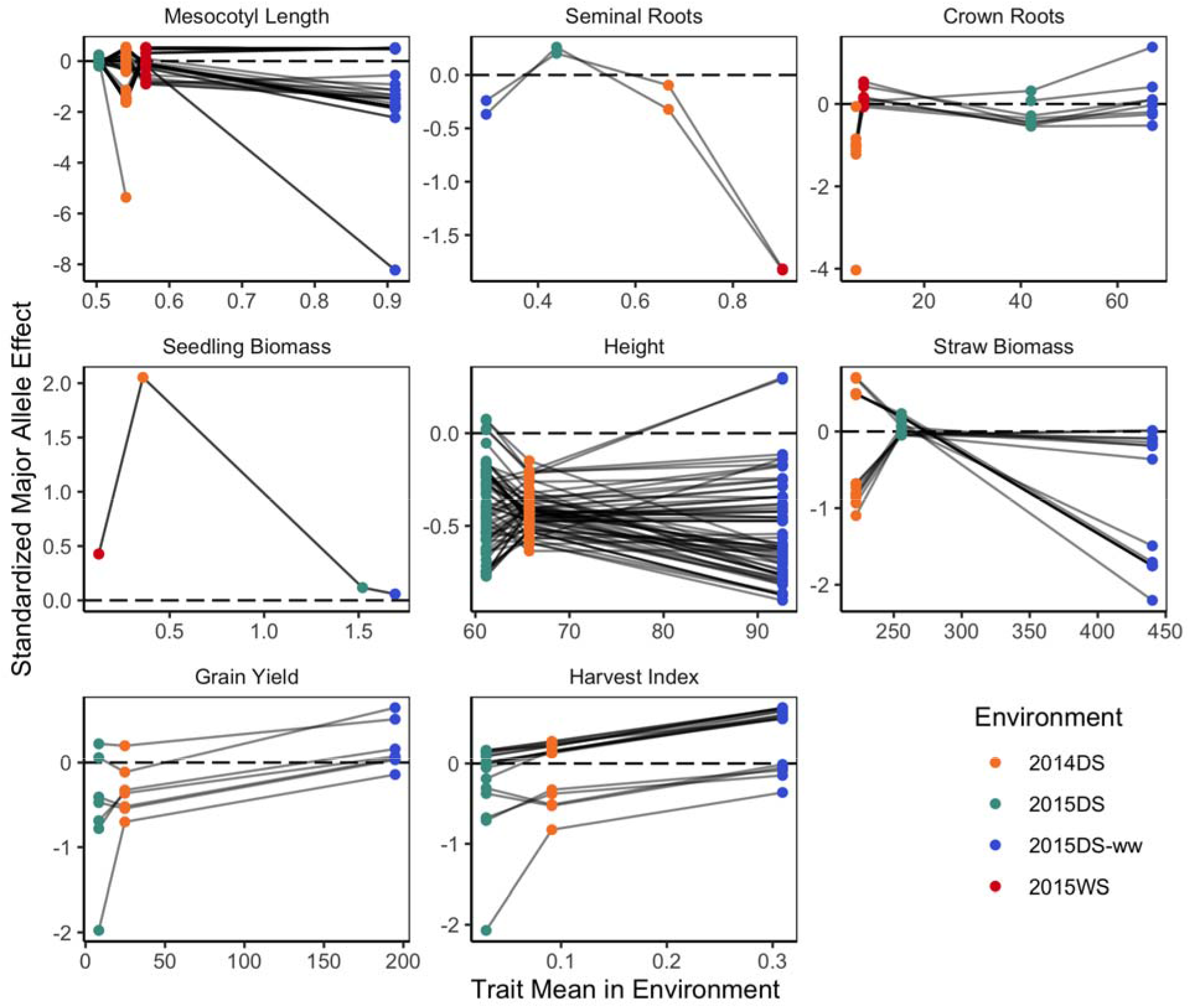
Most genotype by environment allelic effects are conditionally neutral. Major allele trait effects standardized to trait standard deviation in each environment (y-axis) for all SNPs with a permuted *p*-value of < 0.1 in at least one environment for that trait, plotted against the trait mean in each environment for the 453 lines common to all four environments (x-axis). Standardized effects of each SNP across environments are connected by gray lines, and the dashed black horizontal line represents effect = 0. Total axial roots not shown: the panel is highly similar to crown roots.

Most allelic effects we observe are conditionally neutral, with large effects in either the well-watered or one of the drought environments that decrease towards no effect at the other environmental extreme (**Figure 7**). For example, we observe two strong associations with grain yield and harvest index in 2015DS (p8-2 and p9-1, **Table S9**), the most stressful environment f**or** those traits. These two peaks have a moderate but non-significant effect on grain yield and harvest index in DS2014, and an even smaller effect in DS2015-ww. In contrast, peak 1-5 has the strongest association with grain yield and harvest index in 2015DS-ww and essentially no effect in the dry season drought environments.

In addition to conditional neutrality, we observe a few associations with antagonistic effects, where alleles have a positive effect on one environmental extreme and a negative effect on the other. The most notable is peak 3-2 where the major allele has a strong negative effect on straw biomass in DS2015-ww, little effect in 2015DS, and a moderate but not significant positive effect in the environment with the lowest mean biomass, 2014DS (**Figure 7; Table S9**).

### Seedling architecture and growth under dry-direct seeding

We designed these experiments to examine several hypotheses about rice grown under dry-direct seeding. First, we hypothesized that emergence and establishment would be important contributors to yield under dry-direct seeding. In 2014DS neither emergence trait is directly predictive of grain yield. However, later first emergence and earlier full emergence are each moderate predictors of increased straw biomass, which in turn predicts increased grain yield (**Figure 4, Table S8**). However, first and full emergence are strongly positively phenotypically and genetically correlated, suggesting that it might be challenging to target rapid establishment through opposing selection on first and full emergence. We also do not detect any trait-marker associations for emergence or establishment (**Figure 5, Table S9**).

Second, we hypothesized that maintenance of seedling growth during drought would be mediated by establishment and below-ground architecture, and that higher seedling biomass would translate to better yields. We find that seedling biomass has generally weak phenotypic correlations with any trait except crown root number (**Figure 3, Figure 4**). On the other hand, we find that seedling biomass is strongly positively genetically correlated with crown root number (r = 0.77), negatively genetically correlated with emergence, seminal root number and flowering time (r = −0.11 to −0.19), and positively genetically correlated with establishment time, height, straw biomass, grain yield, and harvest index (r = 0.1 to 0.22; **Table 2**). Similar to emergence, however, we observe only one marginal association for seedling biomass on chromosome 3 in 2014DS (**Figure 5, Table S9**).

**Table 2.** Genetic correlations among traits across environments expressed as Pearson’s correlation coefficient *r* (below diagonal) or Spearman’s *rho* (above diagonal), with only correlations significant at p < 0.05 shown. Note that the strength and direction of the correlations are consistent between *r* and *rho*.

|  | First Emergence | Full Emergence | Establishment Time | Crown Roots | Seminal Roots | Total Axial Roots | Mesocotyl Length | Seedling Biomass | Days to Flower | Height | Straw Biomass | Grain Yield | Harvest Index |
| --- | --- | --- | --- | --- | --- | --- | --- | --- | --- | --- | --- | --- | --- |
| First Emergence | 1 | 0.57 | -0.53 | -0.19 | – | -0.19 | – | -0.21 | 0.18 | -0.14 | -0.09 | -0.32 | -0.3 |
| Full Emergence | 0.59 | 1 | 0.3 | -0.17 | – | -0.17 | – | -0.17 | 0.14 | – | – | -0.31 | -0.26 |
| Establishment Time | -0.54 | 0.33 | 1 | – | – | – | – | 0.09 | -0.09 | 0.11 | – | – | 0.09 |
| Crown Roots | -0.2 | -0.17 | 0.11 | 1 | -0.08 | 1 | – | 0.78 | – | 0.11 | 0.12 | 0.31 | 0.18 |
| Seminal Roots | – | – | – | -0.12 | 1 | – | 0.48 | -0.09 | – | – | – | – | – |
| Total Axial Roots | -0.2 | -0.17 | 0.11 | 1 | -0.09 | 1 | – | 0.78 | – | 0.11 | 0.12 | 0.31 | 0.18 |
| Mesocotyl Length | – | – | – | 0.09 | 0.31 | 0.1 | 1 | – | – | – | – | – | – |
| Seedling Biomass | -0.19 | -0.15 | 0.1 | 0.77 | -0.11 | 0.77 | – | 1 | -0.12 | 0.16 | 0.16 | 0.28 | 0.19 |
| Days to Flower | 0.18 | 0.16 | – | -0.09 | – | -0.09 | – | -0.14 | 1 | – | 0.19 | -0.16 | -0.49 |
| Height | -0.13 | – | 0.1 | 0.12 | – | 0.12 | – | 0.15 | – | 1 | 0.43 | 0.12 | – |
| Straw Biomass | – | – | – | 0.14 | – | 0.14 | – | 0.12 | 0.14 | 0.33 | 1 | 0.37 | -0.13 |
| Grain Yield | -0.27 | -0.25 | – | 0.25 | – | 0.25 | – | 0.22 | -0.16 | 0.12 | 0.38 | 1 | 0.72 |
| Harvest Index | -0.29 | -0.27 | 0.08 | 0.2 | – | 0.21 | – | 0.2 | -0.49 | – | -0.1 | 0.65 | 1 |

Finally, we hypothesized that increased mesocotyl length would allow for a better distributed root system (that is, more seminal roots and a higher total number of axial roots), and that this would translate to plant growth. Mesocotyl length has been proposed as an essential breeding target of dry-direct seeded rice for this reason (Lee *et al*. 2017; Zhan *et al*. 2019). We see evidence for the first part of our hypothesis: mesocotyl length consistently predicts and is positively genetically correlated with seminal root number (**Figure 4**, **Table 2**). However, we do not observe strong relationships between these traits and later plant growth. Instead, crown root number is strongly positively correlated with seedling biomass across environments, and genetically correlated with most agronomic traits (**Figure 4, Table 2**). We observe three strong GWA peaks for crown and total axial root number in 2014DS and an unlinked marginal association in 2015DS-ww (**Figure 5, Table S9**). Associations for mesocotyl length vary by environment: two peaks in 2014DS, five different peaks in 2015DS-ww, and a marginal association at p4-1 in 2015WS (**Table S9**). Three of these peaks (p3-3, p4-1, and p8-1) are shared among traits: strongly associated with crown and total axial roots in 2014DS and mesocotyl length in 2015DS-ww, which we could call trans-environmental pleiotropy. We observe additional pleiotropy: p4-1 is the strongest association with grain yield in 2015DS-ww, p8-1 contributes to height in 2014DS, and p1–3 is associated with height as well as mesocotyl length in 2014DS and (marginally) harvest index in 2015DS-ww.

## Discussion

Across seasons and soil moisture levels at seedling stage, we see clear evidence for a major role of the environment in shaping trait means and variances both directly and through interaction with genetic variation. To observe these dynamics required substantial effort across multiple field seasons, which may explain why similar work has rarely been undertaken. However, if researchers and breeders want to fully understand and shape trait variance and covariance, data on the complex roles of environment and genotype by environment interactions are essential. Here we explore how these complex interactions could be used in rice breeding to achieve sustainable yield growth into the future.

### Growth under seedling stage drought

Rapid seedling establishment is a breeding target for direct seeded rice (Kumar and Ladha 2011). Despite developmental separation between emergence and agronomic traits, our results suggest that selection for rapid emergence may have an indirect positive effect on maintenance of seedling growth and yield components under seedling stage drought. Genotypes that were quicker to establish and invest more in crown roots were larger as seedlings, even following drought, and also exhibited higher grain yield and harvest indices. We observe these phenotypic relationships even though we largely controlled for weed competition, one of the primary concerns behind targeting rapid establishment in dry-direct seeding. As we note earlier, however, rapid establishment involves more concerted emergence: first and full emergence need to be selected in opposite directions, but in our experiments they are positively genetically correlated (*r* = 0.59). In addition, we did not find any genetic associations with emergence or establishment. It could be that emergence and establishment are highly polygenic traits and our design did not have the power to detect these many smaller effect associations. It is also possible that the relative contribution of additive genetic variance to these traits was low.

Of our hypotheses for seedling growth under dry-direct seeding and seedling stage drought, we primarily find evidence that crown root number in seedlings is genetically controlled and contributes to later growth and production. While mesocotyl length was a strong predictor of seminal root number and seedling biomass, these effects did not consistently translate to later growth. Plants grown under seedling stage drought were generally smaller with more seminal roots, took longer to flower, and produced lower yields than plants in the well-watered environment. However, traits were generally not strongly phenotypically correlated across the three drought stress environments. Other studies of dry-direct seeding observe similar variation in phenotypic correlation within and across seasons (Sandhu *et al*. 2015; Yadav *et al*. 2017; Subedi *et al*. 2019), and studies of drought tend to find strong environmental effects (Venuprasad *et al*. 2007; Wade *et al*. 2015). These patterns suggest that while mean trait values shift somewhat predictably with drought, genotype rankings and trait covariances change less predictably across development and environments, even among environments that might be superficially labeled as similar. This further highlights how important it is for quantitative genetic research to study drivers of phenotypic variation across environments: genetic architectures are rarely stable.

### Seasonal differences

Dry season (December–May) at IRRI and in the province of Laguna, Philippines is characterized by lower rainfall and lower weather variability than wet season (June–November) (Sander *et al*. 2017). The differences in rainfall between wet and dry seasons have increased with climate change and are projected to continue increasing, with wet seasons growing wetter and dry seasons drier (Chou *et al*. 2013). Both of these changes reduce yields (Lansigan *et al*. 2000; Roberts *et al*. 2009). Rice is grown in the Philippines during both seasons, with lower area (DS 1.87 versus WS 2.63 million hectares; (Sander *et al*. 2017)) during the dry season but higher yields, most likely due to lower solar radiation (Yoshida 1981; Laza *et al*. 2003; Yang *et al*. 2008). Drought is unsurprisingly a larger concern during the dry season—this is one reason for our emphasis on the dry season—but can still occur during the wet season depending on the sowing date. We see a larger role for environmental variation than genetic variation in controlling trait variance during the wet season, with only two weak marker-trait associations for any trait in our wet season data and generally lower heritabilities. This may be an artefact of field phenotyping constraints: we observe fewer associations for early developmental traits than for agronomic traits across all environments, and we did not collect data on most agronomic traits in our wet season environment due to poor plant growth later in the season.

### sd1/qDTY1.1 *has strong consistent effects in the dry season*

The only genomic region with consistent effects on trait variance is on chromosome 1 near 38.5 Mb at peak p1-4 (**Figure 5, Table S9**). The major allele in this region was associated with reduced plant height in all three dry season environments, but only associated with increased grain yield and harvest index in the well-watered environment (**Figure 6, Table S9**). It was also associated with reduced biomass at harvest in 2014DS. *SD1* or gibberellin 20-oxidase, a key enzyme in the gibberellin biosynthesis pathway, is located at 38.38 Mb in the Nipponbare genome assembly, directly amidst the observed strong associations with height (**Figure 6**). The major green revolution allele *semi-dwarf1* is a loss of function allele for *SD1* that creates shorter plants with higher harvest indices (Angira *et al*. 2019). From resequencing data analyzed in Angira *et al*. (2019), the Global MAGIC parents include three functional *SD1* haplotypes: common haplotype 2 (common in the aromatic and *japonica* subpopulations), common haplotype 4 (common in the *aus* subpopulation), and one rare haplotype in IR 4630-22-2-5-1-3 (**Table S1**). While our GBS dataset has no variants within *SD1*, we observe five haplotypes in the 24 SNPs within 300 kb: the major haplotype shared by eleven parents (with one mutation in Cypress) that is likely *sd1;* two similar haplotypes in Jinbubyeo/Inia Tacuari and IET 14720 that are likely linked to *SD1* haplotype 2; and two divergent haplotypes in IAC 165 and IR 4630-22-2-5-1-3 that are likely linked to *SD1* haplotype 4 and the rare *SD1* haplotype, respectively.

A negative effect locus for yield under drought, *qDTY1.1*, is tightly linked to the *sd1* haplotype at ~38.4–38.9 Mb (Vikram *et al*. 2015), which may explain why we saw a positive effect of this locus on grain yield and harvest index in the well-watered environment but not in any of the drought environments. *qDTY1.1* was characterized under puddled, transplanted, reproductive stage drought, but our observations suggest that it may also play a role in dry-direct seeded rice under seedling stage drought. The strongest relationship with height in both dry season drought environments is within the *qDTY1.1* region, while in the well-watered environment it is with the marker closest to *sd1* (S1_38401520, **Figure 6; Table S9**). Disentangling the composite effect of *sd1* and *qDTY1.1* due to tight physical linkage is outside of the bounds of the Global MAGIC population, at least as currently genotyped, but is a major target for breeders seeking semi-dwarf, high yielding under drought varieties.

### Digging in to genetic roots

Our association peak on chromosome 4 (p4-1) was pleiotropically associated with crown and total axial root number, mesocotyl length, and grain yield across different environments. This peak overlaps with many previously discovered QTL for both yield components (number of panicles, grains per panicle, grain weight) and root traits (root number, thickness, depth, length, ratio of deep rooting, root angle, and pulling force), including strong associations with the number of nodal roots in two seasons of dry-direct seeding in a five-parent *aus-indica* MAGIC population (Subedi *et al*. 2019; Sandhu *et al*. 2019a) and the well-characterized *DRO2* locus (Uga *et al*. 2013; Kitomi *et al*. 2015). Of the more than 1000 annotated genes within p4-1, there are at least four genes with known function in root development: the glutamate receptor–like gene *GLR3.1* involved in maintaining apical root meristem development in seedlings (Os04g0585200; (Li *et al*. 2006), *NARROW LEAF1* (*NAL1*, Os04g0615000), which regulates adventitious root development in an auxin-mediated way (Cho *et al*. 2014), diacylglycerol kinase *OsDGK1* that acts to modulate root architecture (Os04g0634700; (Yuan *et al*. 2019), and the jasmonic acid-induced putative receptor-like gene *OsRLK* (Os04g0659300; (Jiang *et al*. 2007). *NAL1*, also known as *SPIKE*, has pleiotropic effects on multiple plant architecture traits and yield components and has been under strong selection by breeders (Takai *et al*. 2013; Fujita *et al*. 2013), making the gene an attractive candidate to underlie variation in mesocotyl length, crown root number, and grain yield here. We note, however, that our p4-1 associations may represent the causal effects of multiple linked loci rather than the pleiotropic effect of one locus.

Two additional peaks were associated with crown and total axial roots in 2014DS and with mesocotyl length in 2015DS-ww: p3-3 and p8-1. Peak 8-1 was also associated with height in 2014DS. These loci, too, overlap with those from previous research: p8-1 with a meta-QTL the ratio of deep rooting under water deficit (Khahani *et al*. 2021). Peak 3-3 overlaps with the major QTL for mesocotyl length in seedlings qMel-3 (Wu *et al*. 2015; Lee *et al*. 2017; Zhan *et al*. 2019), as well as a meta-QTL for root number (Courtois *et al*. 2009). The p3-3 marker strongly associated with crown root number, S3_21631936, sits in the third intron of LOC_Os03g38940.1, an expressed protein of unknown function. The minor alleles at this site and all markers associated with mesocotyl length at p3-3 were donated by the sole Basmati group parent IET 14720, and increased mesocotyl length by 1.8 standard deviations in 2015DS-ww and crown root number by more than four standard deviations in 2014DS.

Peak p8-2 was associated with drought response index, grain yield, and harvest index in 2015DS and overlaps with a QTL hot spot for deep roots and yield across a range of drought-prone environments (Wade *et al*. 2015). The Global MAGIC lines with this minor allele would be good candidates for further characterization of root architecture and yield under drought stress.

### What to select for when most alleles are conditionally neutral

Where the genetic architectures of important traits are highly dependent on environmental interactions, there is likely no single genetic solution to increasing yields. For selective breeding to succeed in generating varieties with stable traits, targeted alleles need to either act consistently across environments (constitutively) or have a positive effect in some environments with no negative effect in others (conditional neutrality). If alleles exhibit environmental antagonism, with positive effects in some environments and negative in others, then environmentally targeted breeding is necessary. Fortunately, antagonistic effect alleles are rare in our observations, although not as rare as alleles with constitutive effects. One such locus is peak p3-2, where the major haplotype reduced straw biomass by ~1.75 standard deviations in 2015DS-ww and had a positive (although non-significant) effect of ~0.5 standard deviations at the other environmental extreme (2014DS). This region overlaps with several previously characterized QTL for shoot length and biomass, and one potential candidate gene in this region is *OsAGO7* (Os03g0449200), which has pleiotropic effects on plant architecture via regulation of the shoot apical meristem and leaf development (Itoh *et al*. 2000; Shi *et al*. 2007).

The vast majority of the allelic effects we observe are conditionally neutral, which means that to detect them required that we evaluate our population across multiple environments. This requirement increases the practical investment needed for genetic discovery, but may prove essential for increasing our rate of yield improvement. Pyramiding of conditionally neutral positive-effect alleles would enable the creation of varieties with high trait stability. We explore one such possibility for grain yield in our data. In the most stressful environment for yield (2015DS), the minor allele at marker S9_11320388 (p9-1) increased grain yield and harvest index by approximately two standard deviations (31 g m^-2^ or HI of 0.1). This allele had a weaker but still positive effect in 2014DS (~0.75 SD), and little effect in 2015DS-ww. Peak 9-1 overlaps with a previously characterized QTL for leaf relative water content under drought (Price *et al*. 2002). S9_11320388 is in the first intron of the leucoanthocyanidin dioxygenase-like gene Os09t0354100. The minor allele at S9_11320388 is a nonsynonymous G–C SNP that changes residue 32 from aspartic acid to histidine in the protein structure of Os09t0354100, although we note that likely this marker is not causal but only linked to the causal variant for p9-1. Twentyeight genotyped MAGIC lines inherited the minor allele from parent IR77186-122-2-2-3, of which we phenotyped ten.

In contrast, the p4-1 major allele at S4_32171832 increased grain yields by ~0.64 SD or 78 g m^-2^ in 2015DS-ww, with essentially no effect in either dry season drought environment. Parents Cypress, Fedearroz 50, Inia Tacuari, IET 14720, and Jinbubyeo have the minor allele at this site, as does IR77186-122-2-2-3, such that the positive two-locus genotype of S4_32171832:major with S9_11320388:minor is not found in any parent genotype. We do see this two-locus genotype in at least six genotyped MAGIC lines (MG-7277, MG-7278, MG-7279, MG-7362, MG-7544, MG-7646), two of which we phenotyped (MG-7279 & MG-7544). There are also several MAGIC lines with missing genotypes at one of these loci which could have the positive combination, including MG-7542, which had the lowest yield stability index of any line. In fact, the mean YSI for MAGIC lines carrying S4_32171832:major or S9_11320388:minor, either in combination or with a missing genotype at the other locus, is significantly lower or more stable than would be expected at random (permutation test, *p* = 0.004). This example illustrates the potential of pyramiding positive effect alleles that are conditionally neutral in alternate environments.

### Conclusions

Our results highlight two key points: (1) complex recombinant populations like the Global MAGIC have great potential for generation of multiallelic recombinants with transgressive trait values, (2) in using them it is important to generate as many recombinant lines as possible, especially in a primarily selfing crop like rice where much variation is at low frequency. The likelihood of any specific combination of rare alleles in a MAGIC population at unlinked loci is approximately 1/N^L, where N is the number of parent varieties and L is the number of loci. Here again practical consideration for space, time, and labor must be balanced against the potential for discovery: even in a 1200 line 16-parent MAGIC population like the Global MAGIC, which required considerable investment to create, the likelihood of a two-locus rare allele recombinant genotype is 1/256, approximately 0.4% or 4–5 lines. In addition, our observation of predominant conditional neutrality among allelic effects emphasizes the importance of multi-location testing to identify the most promising lines in a complex population like the Global MAGIC, but at the same time implies that favorable alleles in some environments would likely show few detrimental effects in other environments.

## Acknowledgements

We would like to acknowledge R. Torres, C. Cabral, L. Holongbayan, M. Quintana, E. Mico, L. Navarro, A. Reyes, and N. Driz for technical support, and the Global Rice Science Partnership (GRiSP) for funding the field trials at IRRI. BTM was supported in part by a NSF National Plant Genome Initiative Postdoctoral Fellowship (award #1523752) and by the University of Massachusetts Boston.

## Supplemental Figure and Table Legends

**Figure S1.** Soil water potentials (mean -kPa ± SD across 9–12 tensiometers) at 15 (top) and 30 cm (bottom) depth for each of the drought stress environments. Only measurements at 30 cm were taken in 2015DS. Soil water potentials were only monitored during drought in 2014DS as the plot was maintained well-watered afterwards, while the other environments were monitored for longer as stress was resumed (2015DS) or remained mild (2015WS).

**Figure S2**. Alignments of sequences from which the genotypes in this study are derived against two genomes, (ssp. *japonica* Nipponbarre assembly IRGSP-1.0/MSU7 and ssp. *indica* 93-11 assembly ASM465v1). An additional set of sequences aligned against only one genome (18% of sites in the Nipponbarre alignment, 16% in the 93-11 alignment). See also **Table S4**.

**Figure S3**. Heatmap of the kinship matrix estimated using the Centered Identity-by-State method in TASSEL 5 for 1184 Global MAGIC population S6 recombinant inbred lines. Higher values indicate higher relatedness. Mean kinship = −0.001 ± 0.106 SD. Note the essentially flat population structure.

**Figure S4.** Trait distributions for Global MAGIC lines (turquoise) and parents (orange) for all traits in the four environments. Boxplots (center line: median, box edges: 25^th^ and 75^th^ percentiles, whiskers: ~95% CI) are plotted over observed values (points).

**Figure S5**. Histograms of days to flower for all four environments. The lines that flowered earlier than 50 days in 2015WS are highlighted in red.

**Figure S6**. Allele frequencies in 1176 S6 generation MAGIC lines versus allele frequencies in the subset of 566 MAGIC lines phenotyped in this study at the 7795 variants aligned against the Nipponbare genome.

**Figure S7**. Compared to **Figure 7**, here effects are in trait rather than variance units. Major allele trait effects (y-axis) for all SNPs with a permuted *p*-value of < 0.1 in at least one environment for that trait, plotted against the trait mean in each environment for the 453 lines grown in all four environments (x-axis). Effects of each SNP across environments are connected by gray lines, and the dashed black horizontal line represents effect = 0. Total axial roots not shown: the panel is highly similar to crown roots.

**Table S1**. Passport and other relevant information for the 16 parent lines of the rice Global MAGIC population created at the International Rice Research Institute, including information for lines sequenced as part of the 3K rice genomes project with the same variety names. Data from: (Bandillo *et al*. 2013; 3,000 rice genomes project 2014; Angira *et al*. 2019).

**Table S2.** Experimental entries across all four environments. Checks and Parent lines were replicated six times in 2014DS, four times in both 2015DS environments, and three times in 2015WS in augmented designs.

**Table S3.** Model fit parameters for the structural equation models built within each environment (and with subsets of traits for 2014DS to match other environments).

**Table S4.** Alignment and SNP calling statistics of our GBS TASSEL tag database against two rice genomes (ssp. *japonica* Nipponbare assembly IRGSP-1.0/MSU7 and ssp. *indica* 93-11 assembly ASM465v1).

**Table S5.** Proportion of MAGIC lines with transgressive trait values and summary statistics for one-sided Ansari-Bradley tests that variance of the Global MAGIC lines (N = 236–538, depending on trait and environment) is greater than variance of the parent lines (N = 16) for all measured traits in the four environments. Note that unequal sample sizes greatly reduce the power of this test to reject the null hypothesis of no difference, so only large differences in variance are likely to be statistically significantly different.

**Table S6.** Eigenvalues and percentage of individual variance explained (PVE) by the top 12 principal components (PCs) of phenotypic variance across environments. We examine in more detail PCs with eigenvalues > 1 in **Figure 2 and Table S7**.

**Table S7.** Correlations of each trait with the first four principal components of phenotypic variance across environments. Only correlations with *p* < 0.05 are shown.

**Table S8.** Structural equation model estimates for all parameters within each environment, including trait variances and intercepts, trait covariances, and trait predictions.

**Table S9.** Genomic loci associated with trait variance across environments. Each peak is defined by the span of associations with permuted *p* < 0.1 that are ≤ 2 Mb from the next associated marker (average breakdown of LD). Major peaks with at least one association with permuted *p* < 0.05 are named pChromosome–CountofPeaksAlongChromosome, while associations with 0.05 < permuted *p* < 0.1 are named pChromosome–minor. The column Markers denotes the number of associations in the range of Position on that chromosome. The following columns give additional details for the strongest marker-trait association for that peak in that environment.

## References

3,000 rice genomes project, 2014 The 3,000 rice genomes project. Gigascience 3: 7.

Anderson, J. T., J. H. Willis, and T. Mitchell-Olds, 2011 Evolutionary genetics of plant adaptation. Trends Genet. 27: 258–266.

Angira, B., C. K. Addison, T. Cerioli, D. B. Rebong, D. R. Wang et al., 2019 Haplotype Characterization of the sd1 Semidwarf Gene in United States Rice. Plant Genome 12: 190010.

Balwinder-Singh, A. J. McDonald, V. Kumar, S. P. Poonia, A. K. Srivastava et al., 2019 Taking the climate risk out of transplanted and direct seeded rice: Insights from dynamic simulation in Eastern India. Field Crops Res. 239: 92–103.

Bandillo, N., C. Raghavan, P. A. Muyco, M. A. L. Sevilla, I. T. Lobina et al., 2013 Multi-parent advanced generation inter-cross (MAGIC) populations in rice: progress and potential for genetics research and breeding. Rice 6: 11.

Bates, D., M. Mächler, B. Bolker, and S. Walker, 2015 Fitting Linear Mixed-Effects Models Using lme4. Journal of Statistical Software, Articles 67: 1–48.

Becker, H. C., and J. Leon, 1988 Stability analysis in plant breeding. Plant Breed. 101: 1–23.

Bidinger, F. R., V. Mahalakshmi, and G. D. P. Rao, 1987 Assessment of drought resistance in pearl millet (Pennisetum americanum (L.) Leeke). II. Estimation of genotype response to stress. Aust. J. Agric. Res. 38: 49–59.

Bossa-Castro, A. M., C. Tekete, C. Raghavan, E. E. Delorean, A. Dereeper et al., 2018 Allelic variation for broad-spectrum resistance and susceptibility to bacterial pathogens identified in a rice MAGIC population. Plant Biotechnol. J.

Chou, C., J. C. H. Chiang, C.-W. Lan, C.-H. Chung, Y.-C. Liao et al., 2013 Increase in the range between wet and dry season precipitation. Nat. Geosci. 6: 263–267.

Cho, S.-H., S.-C. Yoo, H. Zhang, J.-H. Lim, and N.-C. Paek, 2014 Rice NARROW LEAF1 Regulates Leaf and Adventitious Root Development. Plant Mol. Biol. Rep. 32: 270–281.

Conner, J. K., and D. L. Hartl, 2004 A primer of ecological genetics. Sinauer Associates Incorporated.

Core Team, R., and Others, 2013 R: a language and environment for statistical computing. R Foundation for statistical computing, Vienna.

Courtois, B., N. Ahmadi, F. Khowaja, A. H. Price, J.-F. Rami et al., 2009 Rice Root Genetic Architecture: Meta-analysis from a Drought QTL Database. Rice 2: 115–128.

De Mendiburu, F., and R. Simon, 2015 Agricolae - Ten years of an open source statistical tool for experiments in breeding, agriculture and biology: PeerJ PrePrints e1748.

Descalsota, G. I. L., B. P. M. Swamy, H. Zaw, M. A. Inabangan-Asilo, A. Amparado et al., 2018 Genome-Wide Association Mapping in a Rice MAGIC Plus Population Detects QTLs and Genes Useful for Biofortification. Front. Plant Sci. 9: 1347.

Devkota, K. P., Sudhir-Yadav, C. M. Khanda, S. J. Beebout, B. K. Mohapatra et al., 2020 Assessing alternative crop establishment methods with a sustainability lens in rice production systems of Eastern India. J. Clean. Prod. 244: 118835.

Dobermann, A., C. Witt, D. Dawe, S. Abdulrachman, H. C. Gines et al., 2002 Site-specific nutrient management for intensive rice cropping systems in Asia. Field Crops Res. 74: 37–66.

Endelman, J. B., and J.-L. Jannink, 2012 Shrinkage estimation of the realized relationship matrix. G3 2: 1405–1413.

Fujita, D., K. R. Trijatmiko, A. G. Tagle, M. V. Sapasap, Y. Koide et al., 2013 NAL1 allele from a rice landrace greatly increases yield in modern indica cultivars. Proc. Natl. Acad. Sci. U. S. A. 110: 20431–20436.

Fukai, S., and M. Ouk, 2013 Increased productivity of rainfed lowland rice cropping systems of the Mekong region. Crop Pasture Sci. 63: 944–973.

Glaubitz, J. C., T. M. Casstevens, F. Lu, J. Harriman, R. J. Elshire et al., 2014 TASSEL-GBS: a high capacity genotyping by sequencing analysis pipeline. PLoS One 9: e90346.

Haefele, S. M., Y. Kato, and S. Singh, 2016 Climate ready rice: Augmenting drought tolerance with best management practices. Field Crops Res. 190: 60–69.

Ismail, A. M., and G. Atlin, 2019 Incorporating stress tolerance in rice, pp. 224 in Sustaining Global Food Security: The Nexus of Science and Policy, edited by R. Zeigler. Csiro Publishing.

Itoh, J. I., H. Kitano, M. Matsuoka, and Y. Nagato, 2000 Shoot organization genes regulate shoot apical meristem organization and the pattern of leaf primordium initiation in rice. Plant Cell 12:2161–2174.

Jiang, J., J. Li, Y. Xu, Y. Han, Y. Bai et al., 2007 RNAi knockdown of Oryza sativa root meander curling gene led to altered root development and coiling which were mediated by jasmonic acid signalling in rice. Plant Cell Environ. 30: 690–699.

Josse, J., F. Husson, and Others, 2016 missMDA: a package for handling missing values in multivariate data analysis. J. Stat. Softw. 70: 1–31.

Kawahara, Y., M. de la Bastide, J. P. Hamilton, H. Kanamori, W. R. McCombie et al., 2013 Improvement of the Oryza sativa Nipponbare reference genome using next generation sequence and optical map data. Rice 6: 4.

Khahani, B., E. Tavakol, V. Shariati, and L. Rossini, 2021 Meta-QTL and ortho-MQTL analyses identified genomic regions controlling rice yield, yield-related traits and root architecture under water deficit conditions. Sci. Rep. 11: 6942.

Kitomi, Y., N. Kanno, S. Kawai, T. Mizubayashi, S. Fukuoka et al., 2015 QTLs underlying natural variation of root growth angle among rice cultivars with the same functional allele of DEEPER ROOTING 1. Rice 8: 16.

Kumar, V., and J. K. Ladha, 2011 Chapter Six - Direct Seeding of Rice: Recent Developments and Future Research Needs, pp. 297–413 in Advances in Agronomy, edited by D. L. Sparks. Academic Press.

Langmead, B., C. Wilks, V. Antonescu, and R. Charles, 2019 Scaling read aligners to hundreds of threads on general-purpose processors. Bioinformatics 35: 421–432.

Lansigan, F. P., W. L. de los Santos, and J. O. Coladilla, 2000 Agronomic impacts of climate variability on rice production in the Philippines. Agric. Ecosyst. Environ. 82: 129–137.

Laza, M. R., S. Peng, S. Akita, and H. Saka, 2003 Contribution of Biomass Partitioning and Translocation to Grain Yield under Sub-Optimum Growing Conditions in Irrigated Rice. Plant Prod. Sci. 6: 28–35.

Lee, H.-S., K. Sasaki, J.-W. Kang, T. Sato, W.-Y. Song et al., 2017 Mesocotyl Elongation is Essential for Seedling Emergence Under Deep-Seeding Condition in Rice. Rice 10: 32.

Lê, S., J. Josse, F. Husson, and Others, 2008 FactoMineR: an R package for multivariate analysis. J. Stat. Softw. 25: 1–18.

Leung, H., C. Raghavan, B. Zhou, R. Oliva, I. R. Choi et al., 2015 Allele mining and enhanced genetic recombination for rice breeding. Rice 8: 34.

Li, J., S. Zhu, X. Song, Y. Shen, H. Chen et al., 2006 A rice glutamate receptor-like gene is critical for the division and survival of individual cells in the root apical meristem. Plant Cell 18: 340–349.

Meng, L., L. Guo, K. Ponce, X. Zhao, and G. Ye, 2016a Characterization of Three Rice Multiparent Advanced Generation Intercross (MAGIC) Populations for Quantitative Trait Loci Identification. Plant Genome 9.:

Meng, L., X. Zhao, K. Ponce, G. Ye, and H. Leung, 2016b QTL mapping for agronomic traits using multi-parent advanced generation inter-cross (MAGIC) populations derived from diverse elite indica rice lines. Field Crops Res. 189: 19–42.

Neue, H.-U., 1993 Methane Emission from Rice Fields. Bioscience 43: 466–474.

Ogawa, D., Y. Nonoue, H. Tsunematsu, N. Kanno, T. Yamamoto et al., 2018a Discovery of QTL Alleles for Grain Shape in the Japan-MAGIC Rice Population Using Haplotype Information. G3 8: 3559–3565.

Ogawa, D., E. Yamamoto, T. Ohtani, N. Kanno, H. Tsunematsu et al., 2018b Haplotype-based allele mining in the Japan-MAGIC rice population. Sci. Rep. 8: 4379.

Pandey, S., M. Mortimer, L. Wade, T. P. Tuong, K. Lopez et al. (Eds.), 2002 Direct Seeding: Research Strategies and Opportunities. International Rice Research Institute.

Price, A. H., J. Townend, M. P. Jones, A. Audebert, and B. Courtois, 2002 Mapping QTLs associated with drought avoidance in upland rice grown in the Philippines and West Africa. Plant Mol. Biol. 48: 683–695.

Raghavan, C., R. Mauleon, V. Lacorte, M. Jubay, H. Zaw et al., 2017 Approaches in Characterizing Genetic Structure and Mapping in a Rice Multiparental Population. G3 7: 1721–1730.

Ray, D. K., N. D. Mueller, P. C. West, and J. A. Foley, 2013 Yield Trends Are Insufficient to Double Global Crop Production by 2050. PLoS One 8: e66428.

Roberts, M. G., D. Dawe, W. P. Falcon, and R. L. Naylor, 2009 El Niño–Southern Oscillation Impacts on Rice Production in Luzon, the Philippines. J. Appl. Meteorol. Climatol. 48: 1718–1724.

Rosseel, Y., 2012 Lavaan: An R package for structural equation modeling and more. Version 0.5--12 (BETA). J. Stat. Softw. 48: 1–36.

Sakai, H., S. S. Lee, T. Tanaka, H. Numa, J. Kim et al., 2013 Rice Annotation Project Database (RAP-DB): an integrative and interactive database for rice genomics. Plant Cell Physiol. 54: e6.

Sander, B. O., R. Wassmann, L. K. Palao, and A. Nelson, 2017 Climate-based suitability assessment for alternate wetting and drying water management in the Philippines: a novel approach for mapping methane mitigation potential in rice production. Carbon Management 8: 331–342.

Sandhu, N., S. R. Subedi, V. K. Singh, P. Sinha, S. Kumar et al., 2019a Deciphering the genetic basis of root morphology, nutrient uptake, yield, and yield-related traits in rice under dry direct-seeded cultivation systems. Sci. Rep. 9: 9334.

Sandhu, N., R. O. Torres, M. T. Sta Cruz, P. C. Maturan, R. Jain et al., 2015 Traits and QTLs for development of dry direct-seeded rainfed rice varieties. J. Exp. Bot. 66: 225–244.

Sandhu, N., R. B. Yadaw, B. Chaudhary, H. Prasai, K. Iftekharuddaula et al., 2019b Evaluating the Performance of Rice Genotypes for Improving Yield and Adaptability Under Direct Seeded Aerobic Cultivation Conditions. Front. Plant Sci. 10: 159.

Shi, Z., J. Wang, X. Wan, G. Shen, X. Wang et al., 2007 Over-expression of rice OsAGO7 gene induces upward curling of the leaf blade that enhanced erect-leaf habit. Planta 226: 99–108.

Subedi, S. R., N. Sandhu, V. K. Singh, P. Sinha, S. Kumar et al., 2019 Genome-wide association study reveals significant genomic regions for improving yield, adaptability of rice under dry direct seeded cultivation condition. BMC Genomics 20: 471.

Takai, T., S. Adachi, F. Taguchi-Shiobara, Y. Sanoh-Arai, N. Iwasawa et al., 2013 A natural variant of NAL1, selected in high-yield rice breeding programs, pleiotropically increases photosynthesis rate. Sci. Rep. 3: 2149.

Tilman, D., C. Balzer, J. Hill, and B. L. Befort, 2011 Global food demand and the sustainable intensification of agriculture. Proc. Natl. Acad. Sci. U. S. A. 108: 20260–20264.

Uga, Y., E. Yamamoto, N. Kanno, S. Kawai, T. Mizubayashi et al., 2013 A major QTL controlling deep rooting on rice chromosome 4. Sci. Rep. 3: 3040.

Venuprasad, R., H. R. Lafitte, and G. N. Atlin, 2007 Response to Direct Selection for Grain Yield under Drought Stress in Rice. Crop Sci. 47: 285–293.

Vikram, P., B. P. M. Swamy, S. Dixit, R. Singh, B. P. Singh et al., 2015 Drought susceptibility of modern rice varieties: an effect of linkage of drought tolerance with undesirable traits. Sci. Rep. 5: 14799.

Wade, L. J., V. Bartolome, R. Mauleon, V. D. Vasant, S. M. Prabakar et al., 2015 Environmental Response and Genomic Regions Correlated with Rice Root Growth and Yield under Drought in the OryzaSNP Panel across Multiple Study Systems. PLoS One 10: e0124127.

Wang, W., R. Mauleon, Z. Hu, D. Chebotarov, S. Tai et al., 2018 Genomic variation in 3,010 diverse accessions of Asian cultivated rice. Nature 557: 43–49.

Wang, W., S. Peng, H. Liu, Y. Tao, J. Huang et al., 2017 The possibility of replacing puddled transplanted flooded rice with dry seeded rice in central China: A review. Field Crops Res. 214:310–320.

Ware, D., P. Jaiswal, J. Ni, X. Pan, K. Chang et al., 2002 Gramene: a resource for comparative grass genomics. Nucleic Acids Res. 30: 103–105.

Wassmann, R., H. U. Neue, R. S. Lantin, K. Makarim, N. Chareonsilp et al., 2000 Characterization of methane emissions from rice fields in Asia. II. Differences among irrigated, rainfed, and deepwater rice, pp. 13–22 in Methane Emissions from Major Rice Ecosystems in Asia, edited by R. Wassmann, R. S. Lantin, and H.-U. Neue. Springer Netherlands, Dordrecht.

Wu, J., F. Feng, X. Lian, X. Teng, H. Wei et al., 2015 Genome-wide Association Study (GWAS) of mesocotyl elongation based on re-sequencing approach in rice. BMC Plant Biol. 15: 218.

Yadav, S., U. M. Singh, S. M. Naik, C. Venkateshwarlu, P. J. Ramayya et al., 2017 Molecular Mapping of QTLs Associated with Lodging Resistance in Dry Direct-Seeded Rice (Oryza sativa L.). Front. Plant Sci. 8: 1431.

Yang, W., S. Peng, R. C. Laza, R. M. Visperas, and M. L. Dionisio-Sese, 2008 Yield Gap Analysis between Dry and Wet Season Rice Crop Grown under High-Yielding Management Conditions. Agron. J. 100: 1390–1395.

Yonemaru, J.-I., T. Yamamoto, S. Fukuoka, Y. Uga, K. Hori et al., 2010 Q-TARO: QTL Annotation Rice Online Database. Rice 3: 194–203.

Yoshida, S., 1981 Fundamentals of Rice Crop Science. Int. Rice Res. Inst.

Yuan, S., S.-C. Kim, X. Deng, Y. Hong, and X. Wang, 2019 Diacylglycerol kinase and associated lipid mediators modulate rice root architecture. New Phytol. 223: 261–276.

Zaw, H., C. Raghavan, A. Pocsedio, B. P. M. Swamy, M. L. Jubay et al., 2019 Exploring genetic architecture of grain yield and quality traits in a 16-way indica by japonica rice MAGIC global population. Sci. Rep. 9: 19605.

Zhan, J., X. Lu, H. Liu, Q. Zhao, and G. Ye, 2019 Mesocotyl elongation, an essential trait for dry-seeded rice (Oryza sativa L.): a review of physiological and genetic basis. Planta 251: 27.

Zhao, W., J. Wang, X. He, X. Huang, Y. Jiao et al., 2004 BGI□RIS: an integrated information resource and comparative analysis workbench for rice genomics. Nucleic Acids Res. 32: D377–D382.

