## Supplemental Figures for "Strong genotype by environment interactions in the rice Global MAGIC population across seedling stage drought"

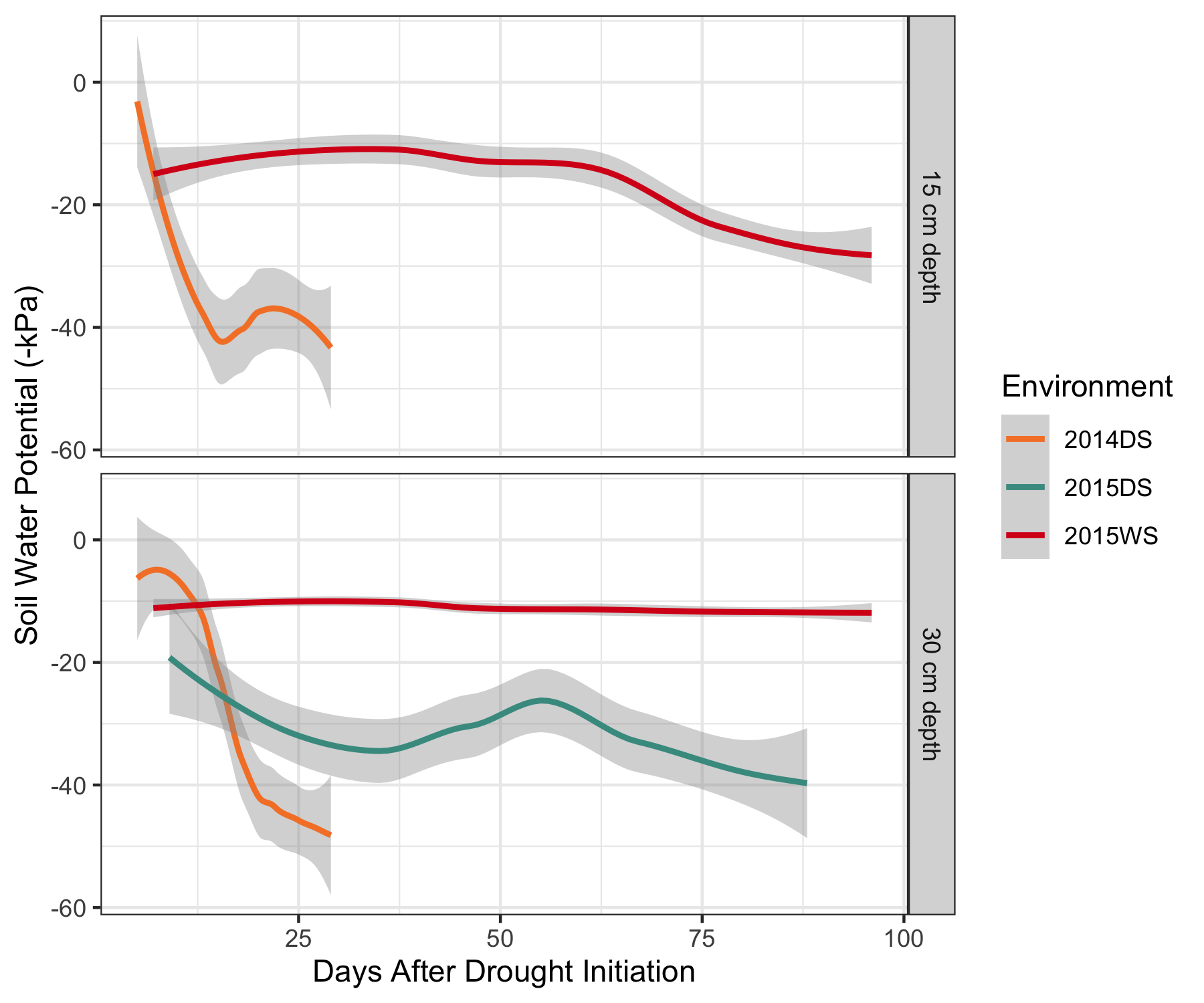

**Figure S1.** Soil water potentials (mean -kPa ± SD across 9–12 tensiometers) at 15 (top) and 30 cm (bottom) depth for each of the drought stress environments. Only measurements at 30 cm were taken in 2015DS. Soil water potentials were only monitored during drought in 2014DS as the plot was maintained well-watered afterwards, while the other environments were monitored for longer as stress was resumed (2015DS) or remained mild (2015WS).

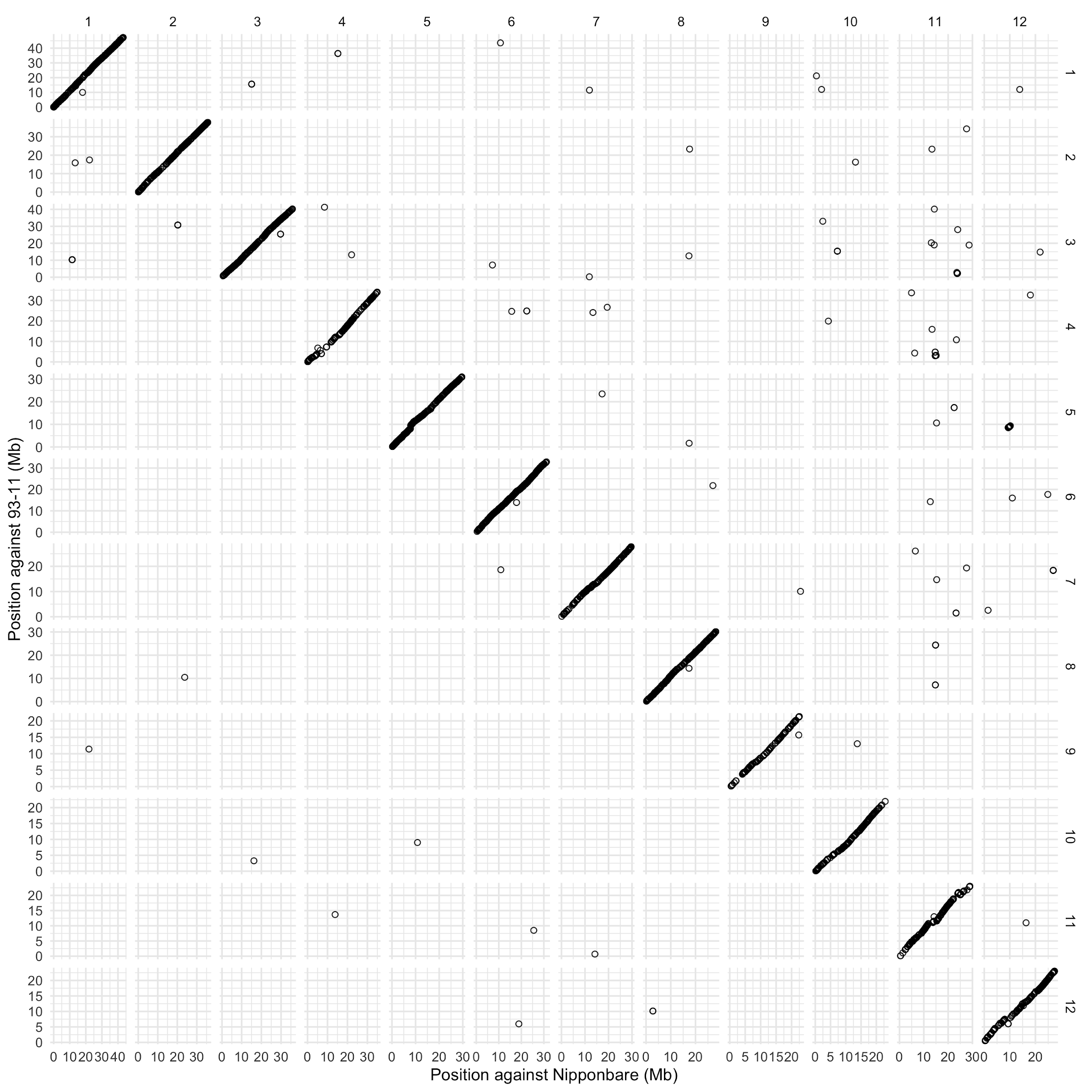

**Figure S2**. Alignments of sequences from which the genotypes in this study are derived against two genomes, (ssp. *japonica* Nipponbarre assembly IRGSP-1.0/MSU7 and ssp. *indica* 93-11 assembly ASM465v1). An additional set of sequences aligned against only one genome (18% of sites in the Nipponbarre alignment, 16% in the 93-11 alignment). See also **Table S4**.

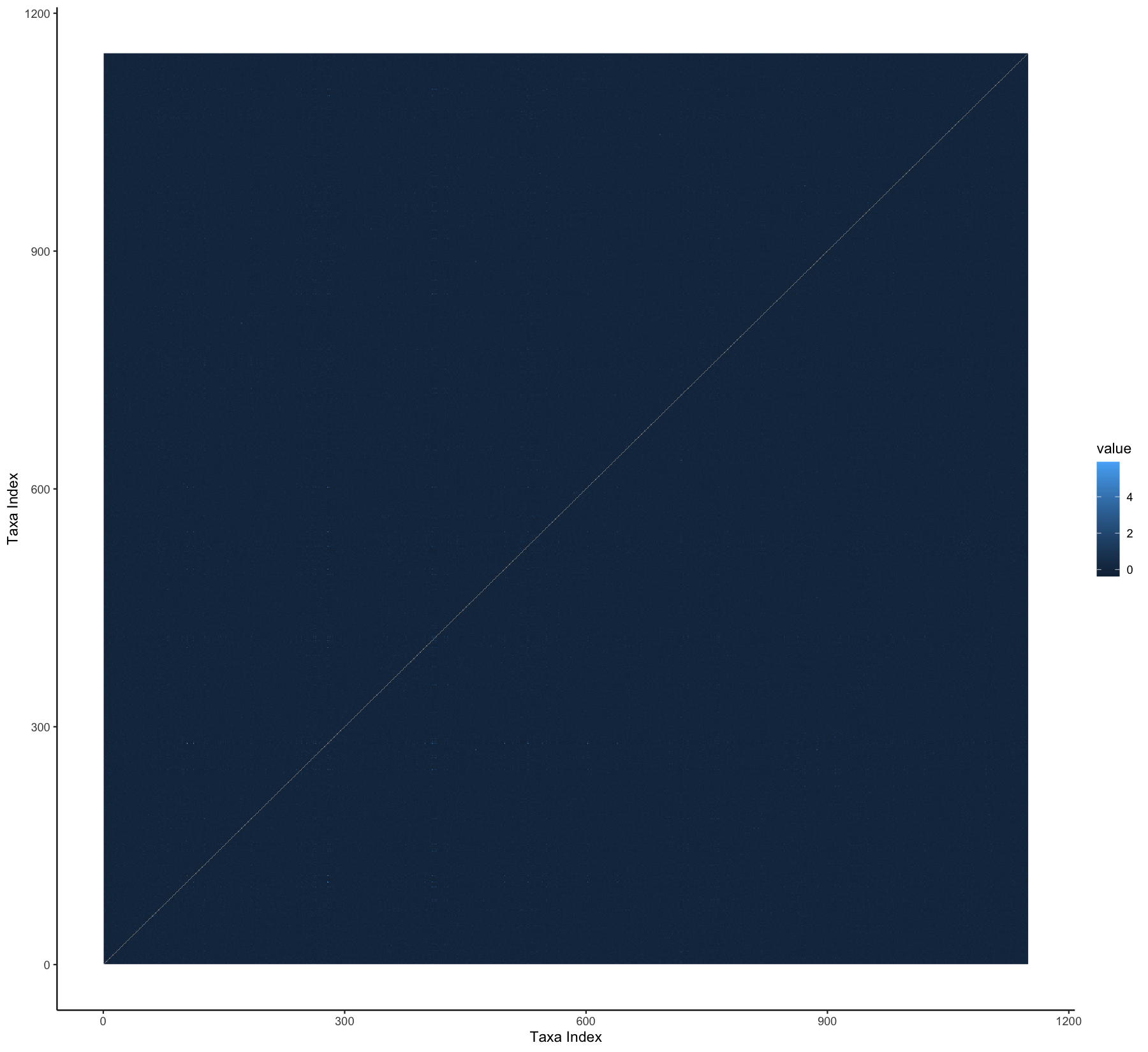

**Figure S3**. Heatmap of the kinship matrix estimated using the Centered Identity-by-State method in TASSEL 5 for 1184 Global MAGIC population S6 recombinant inbred lines. Higher values indicate higher relatedness. Mean kinship = -0.001 ± 0.106 SD. Note the essentially flat population structure.

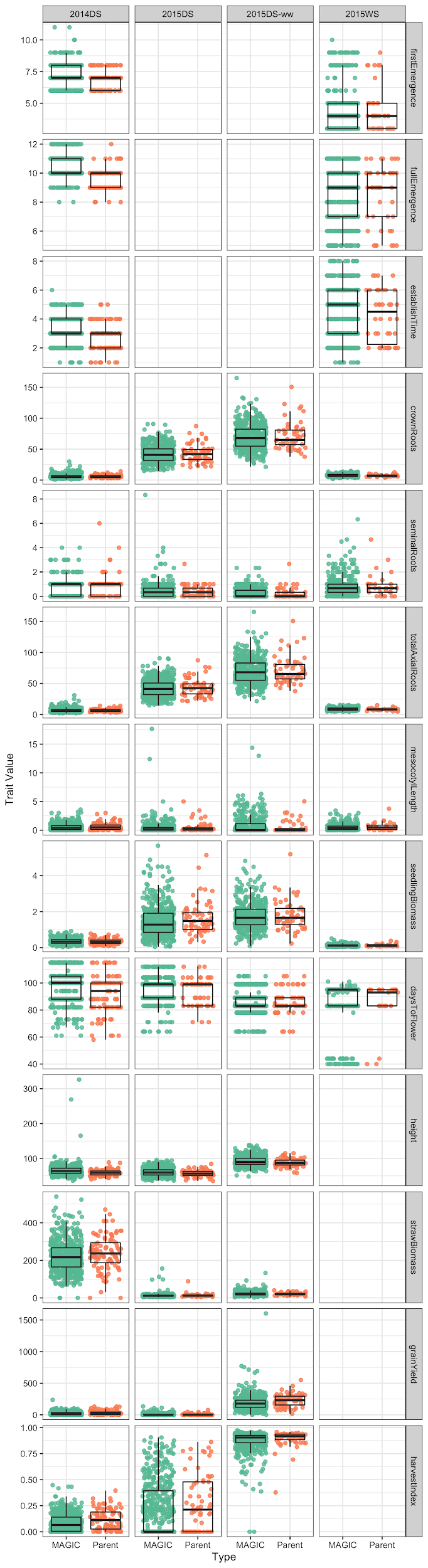

**Figure S4.** Trait distributions for Global MAGIC lines (turquoise) and parents (orange) for all traits in the four environments. Boxplots (center line: median, box edges: 25^th^ and 75^th^ percentiles, whiskers: ~95% CI) are plotted over observed values (points).

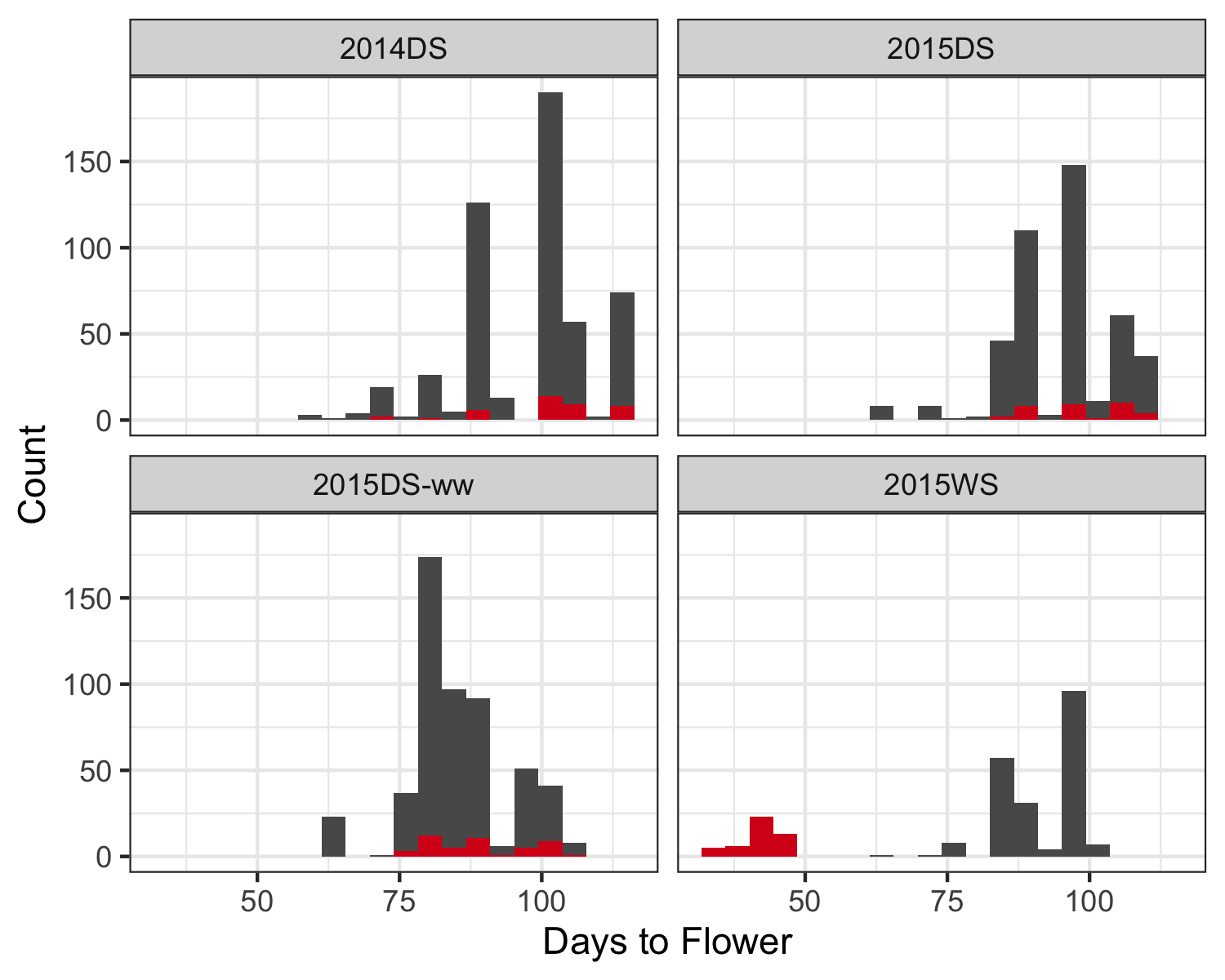

**Figure S5**. Histograms of days to flower for all four environments. The lines that flowered earlier than 50 days in 2015WS are highlighted in red.

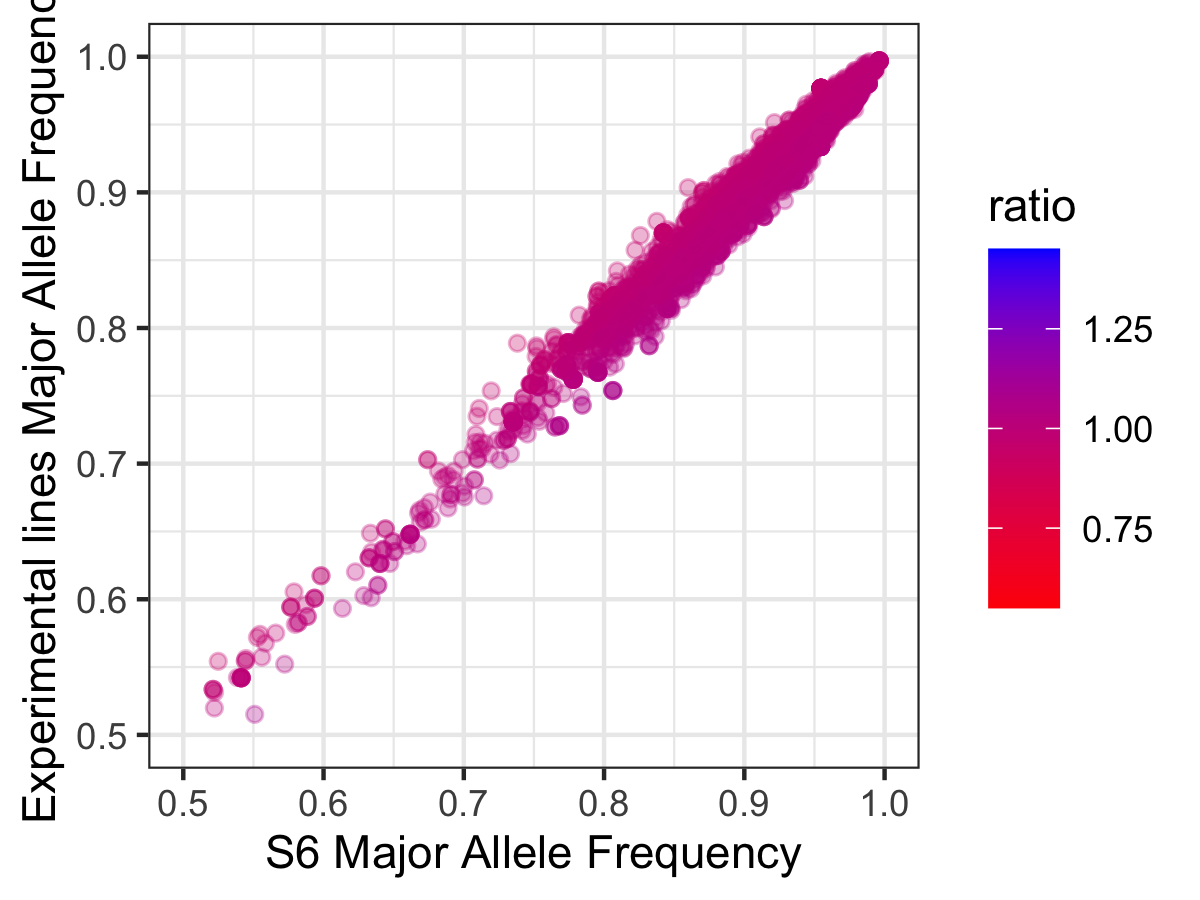

**Figure S6**. Allele frequencies in 1176 S6 generation MAGIC lines versus allele frequencies in the subset of 566 MAGIC lines phenotyped in this study at 7795 variants aligned against the Nipponbare genome.

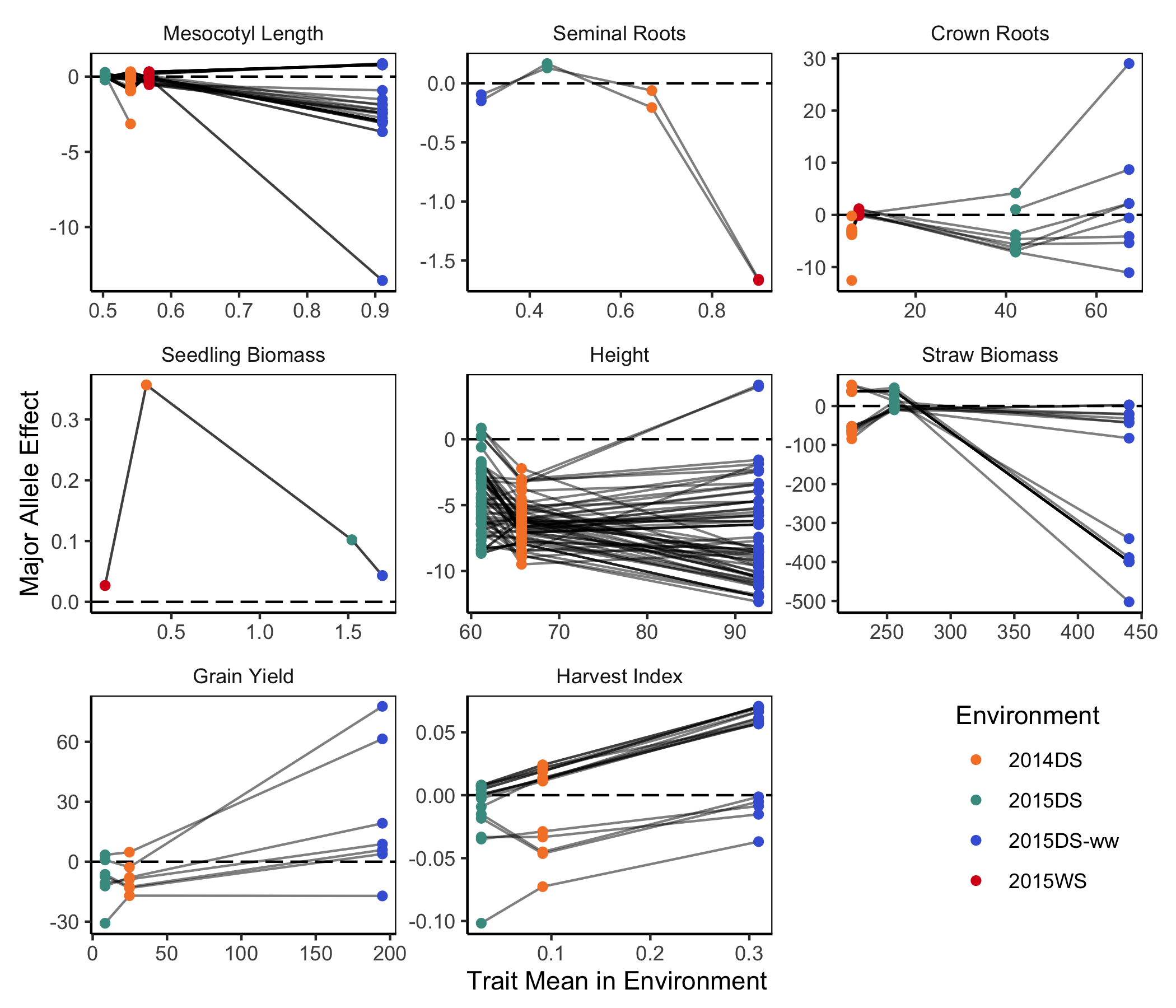

**Figure S7**. Compared to **Figure 7**, here effects are in trait rather than variance units. Major allele trait effects (y-axis) for all SNPs with a permuted *p*-value of < 0.1 in at least one environment for that trait, plotted against the trait mean in each environment for the 453 lines grown in all four environments (x-axis). Effects of each SNP across environments are connected by gray lines, and the dashed black horizontal line represents effect = 0. Total axial roots not shown: the panel is highly similar to crown roots.
